# An adult stem-like cell population generates germline and neurons in the sea anemone *Nematostella vectensis*

**DOI:** 10.1101/2023.01.27.525880

**Authors:** Paula Miramón-Puértolas, Patrick R.H. Steinmetz

**Affiliations:** Michael Sars Centre, University of Bergen, Thormøhlensgt. 55, N-5006 Bergen, Norway

**Author notes:** Corresponding author: Patrick R.H. Steinmetz.

## Abstract

Most genetic research animals (e.g., vertebrates, insects, nematodes) segregate germline and soma during early embryogenesis. In contrast, some highly regenerative bilaterian (e.g., planarians) and non-bilaterian animals (e.g., hydrozoan cnidarians) retain adult stem cells with both germinal and somatic potentials. As these cells have been studied in only few phyla, their biology and evolution remain mostly enigmatic. Here, we aimed to identify and characterize adult stem cells and their cell lineages in the sea anemone *Nematostella vectensis* by combining gene expression analysis, immunostainings, and meganuclease-mediated and CRISPR/Cas9-mediated knock-in reporter lines of conserved germline and multipotency genes (e.g., *vasa2*, *piwi1*). We found a small population of *vasa2*+/*piwi1*+ cells in the gastrodermal folds of juvenile and adult sea anemones that generates germline and a diversity of somatic, mostly proliferative cells. Using a combination of *soxB(2)* neural progenitor and *piwi1* reporter lines, we found that the somatic progeny from *vasa2*+/*piwi1*+ cells includes *soxB(2)*+ neural progenitors. Our results strongly support the existence of an adult Vasa2+/Piwi1+ multipotent stem-like cell population that derives both germline and somatic lineages in *Nematostella*. The similarities of lineages and gene expression profiles between *Nematostella* Vasa2+/Piwi1+ stem-like cells and hydrozoan interstitial stem cells support their evolutionary conservation among cnidarians.

## Introduction

In vertebrates, insects and nematodes, a defined set of primordial germ cells (PGCs) segregates from somatic cell lineages by preformation or induction during early embryogenesis (Extavour and Akam, 2003; Illmensee and Mahowald, 1974; Saitou and Yamaji, 2012; Seervai and Wessel, 2013). PGCs then migrate into the developing gonad and establish a population of germline stem cells (GSCs) that will produce gametes (Richardson and Lehmann, 2010). Notably, animals with embryonic germline segregation exhibit limited regenerative abilities and a lack of asexual reproduction (Rinkevich et al., 2022). In highly regenerative animals (e.g., planarians, acoels, colonial ascidians, hydrozoan cnidarians), a pool of pluri- or multipotent adult stem cells (ASCs) (e.g. planarian neoblasts, sponge archaeocytes) retains both somatic and germinal potential, termed hereafter primordial stem cells (PriSCs) (Solana, 2013), throughout the lifetime of the organism (Baguna et al., 1989; Bode, 1996; David, 2012; Fierro-Constaín et al., 2017; Funayama, 2013; Funayama et al., 2010; Sunanaga et al., 2010). As animals with adult PriSCs are found scattered in few bilaterian phyla and molecular data from non-bilaterian phyla (e.g. sponges, ctenophores, non-hydrozoan cnidarians) is scarce, the evolutionary origin of adult PriSCs remains unclear (Fierro-Constaín et al., 2017; Rinkevich et al., 2022; Sogabe et al., 2019).

In cnidarians, the sister group of bilaterians, germline specification and stem cell biology has almost exclusively been studied in the hydrozoans *Hydra* and *Hydractinia* (Frank et al., 2009). *Hydra* possesses two types of unipotent, epithelial stem cells and a population of PriSCs, the interstitial stem cell population (i-cells) (Frank et al., 2009; Hemmrich et al., 2012; Solana, 2013). The latter consists of a pool of multipotent ASCs that generate both the germline and a set of somatic cell types that includes neurons, gland cells and stinging cells (cnidocytes) (Bosch and David, 1987; Siebert et al., 2019). *Hydractinia* i-cells are pluripotent as they give rise to all lineages, including epithelial cells (Frank et al., 2009; Müller et al., 2004; Varley et al., 2022). As the existence of stem cells in other cnidarian groups (e.g. anthozoans) has remained elusive, i-cells are currently considered as hydrozoan-specific (Gold and Jacobs, 2013; Technau and Steele, 2011).

Anthozoans (i.e., corals and sea anemones), which form the sister group to all other cnidarians, can reproduce both sexually and asexually. Their potential to regenerate an entire body during asexual reproduction or after bisection strongly suggests the presence of ASCs with broad potential (Amiel et al., 2015; Bocharova and Kozevich, 2011; Bythell et al., 2018; Goffredo and Dubinsky, 2018; Reitzel et al., 2007; Röttinger, 2021). The presence of so far unidentified adult PriSCs in anthozoans has been evidenced by recent genome sequencing studies in scleractinian corals, which have identified distinct, post-embryonic single nucleotide variants (SNVs) shared between somatic tissue and the germline of different colony branches (López-Nandam et al., 2023; Vasquez Kuntz et al., 2020).

PriSCs, PGCs and GSCs share the expression of an evolutionarily conserved gene set, the germline multipotency program (GMP) (Juliano et al., 2010). Key components comprise RNA helicases (e.g. *vasa* and *pl10*), RNA-binding (e.g. *tudor*, *nanos*, *pumilio*, *boule* and *bruno*) and RNA-cleaving proteins (e.g. *piwi*) (Alié et al., 2015; Fierro-Constaín et al., 2017; Juliano et al., 2010). They form ribonucleoprotein aggregates (’germline granules’) with target mRNAs that appear as electron-dense fibrillar structures (’nuage’) on an ultrastructural level (Gao and Arkov, 2013; Mahowald, 1962). In the germline of vertebrates, insects and nematodes, these components mediate the suppression of somatic fate by post-transcriptional mRNA regulation and the preservation of genome integrity by silencing transposon activity (Hartung et al., 2014; Robert et al., 2015; Senti and Brennecke, 2010; Zhang et al., 2012). GMP genes have been consistently found expressed in PriSCs of diverse animal species (e.g. neoblasts of planarians and acoels, i-cells of hydrozoan cnidarians, archeocytes and choanocytes of sponges) (De Mulder et al., 2009; Funayama et al., 2010; Rebscher et al., 2008; Reddien et al., 2005; Srivastava et al., 2014; Sunanaga et al., 2010), which suggests that these genes hold conserved roles not only in gametogenesis but also in the maintenance of stemness and multipotency. The coexpression of a subset of GMP genes (e.g. *piwi* and *pumilio*) in vertebrate somatic ASCs (e.g. hematopoietic and neural stem cells) further supports this notion (Sharma et al., 2001; Zhang et al., 2017). GMP genes might thus represent a conserved gene set that allows identifying germline and adult PriSCs, if present, in so far uncharacterized animals (Juliano et al., 2010; Solana, 2013).

Anthozoan gametogenesis has been studied mainly on an ultrastructural level. In *Nematostella vectensis* and other anthozoans, gametes generally develop within the extra-cellular matrix (i.e., mesoglea) of the mesenteries, which are folds of the inner epithelium (i.e., gastrodermis) (Figure 1A) (Eckelbarger et al., 2008; Larkman, 1983; Shikina and Chang, 2016). It is assumed that small oocytes bulge from the surrounding somatic gonad epithelium into the mesoglea, where they continue to grow and mature (Eckelbarger et al., 2008; Moiseeva et al., 2017). The cellular and developmental origin of the germline in anthozoans has recently been addressed by studying the mRNA or protein expression of GMP gene orthologs. In the scleractinian coral *Euphyllia ancora*, Vasa2 and Piwi1 proteins are detected in spermatogonia, oocytes and potential GSCs within the mesenteries (Shikina and Chang, 2016; Shikina et al., 2012; Shikina et al., 2015). Similarly, Vasa2 protein is found in developing gametes and putative GSCs in the mesenteries of juvenile and adult *Nematostella* (Chen et al., 2020; Praher et al., 2017). Orthologs of GMP genes (e.g., *vasa, piwi, tudor*) are expressed broadly during embryonic and early larval development in *Nematostella*. During late larval development, their expression becomes restricted to two clusters of GMP-positive cells located at the oral end of the two primary mesenteries (Chen et al., 2020; Extavour et al., 2005). It has been assumed that these GMP-positive cells constitute PGCs that found a population of GSCs in the mesenteries of *Nematostella* (Chen et al., 2020). However, as the putative somatic potential of these cells has not been addressed, it remains unclear if they consist of PGCs or multipotent adult PriSCs.

**Figure 1.**
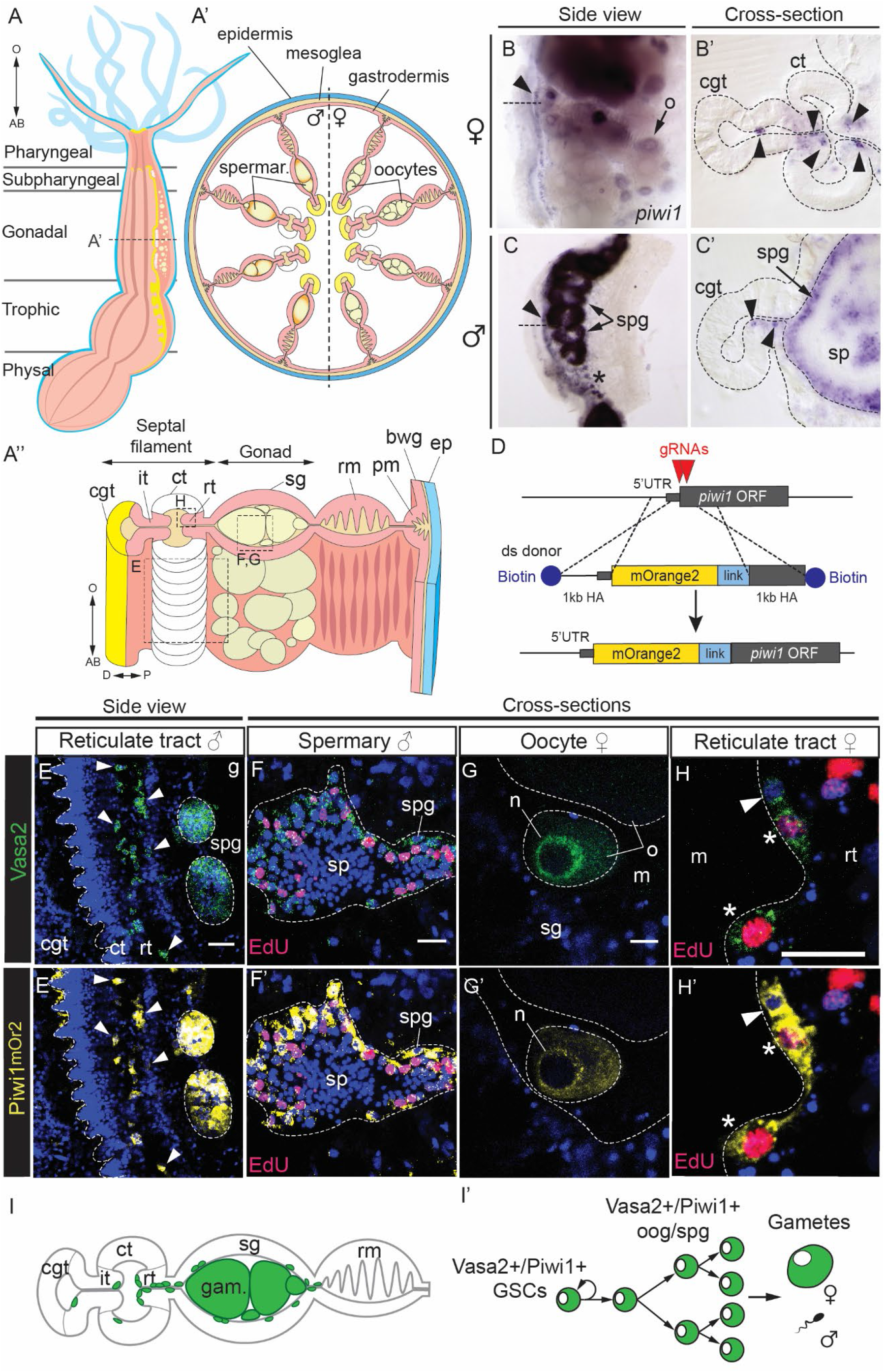
Vasa2 and Piwi1 proteins colocalize to developing gametes and extra-gonadal cells in the mesentery. (A-A’’) Schematics of longitudinal (A) or cross sections (A’, A’’) highlighting the anatomy of adult polyp mesenteries. (A’) and (A’’) represent gonadal mesentery regions. (B-C’) Whole-mount ISH of *piwi1* on dissected gonadal region tissue pieces (B, C) and their cross sections (B’-C’) from female (B, B’) and male (C, C’) mesenteries. Expression was found in the oocytes (o), spermatogonia (spg) and a line of basiepithelial cells at the basis of the septal filament (arrowheads). (D) Schematized strategy for generating a *piwi1^mOr2^* knock-in allele by CRISPR-Cas9-mediated homologous recombination of an mOr2-containing donor construct. (E-H’) Confocal imaging stacks of gonadal mesentery region tissues immunostained for Vasa2 (E-H, green), mOr2-Piwi1 (E’-H’, yellow) and labelled for S-phase by EdU (F, F’, H, H’; 3 days pulse, red) viewed from lateral (E, E’) or at cross section (F-H’) as indicated in (A’’). Vasa2 and mOr2-Piwi1 colocalize to spermatogonia cells (E-F’, spg) and a line of cells along the basis of the septal filament (E, E’, H, H’; white arrowheads). Note that EdU label colocalises to Vasa2+/Piwi1+ peripheric spermatogonia and is rarely found towards centrally located, differentiated, Vasa2-/Piwi1-spermatocytes (sp). Vasa2 and mOr2-Piwi1 colocalize in the nuage (n) of an oocyte (G, G’) and to basiepithelial cells (H, H’; arrowhead), some of which undergo S-phase (H’, asterisks). (I-I’) Schematic summary of basiepithelial Vasa2+/Piwi1+ cells and gametes (I, green) and the potential Vasa2+/Piwi1+ germinal cell lineage (I’). Blue: Hoechst DNA dye. cgt: cnidoglandular tract; it: intermediate tract; ct: ciliated tract; rt: reticulate tract; sg: somatic gonad; rm: retractor muscle; pm: parietal muscle; bwg: body wall gastrodermis; ep: epidermis; o: oocyte; spg: spermatogonia; sp: sperm; sg: somatic gonad, gam: gametes; m: mesoglea. Scale bars: 20μm (E) and 10μm (F-H).

In the present study, we used *in situ* hybridization and Vasa2 protein localization in combination with three transgenic reporter lines to show that Vasa2+/Piwi1+ cells constitute a population of adult stem-like cells that generates gametes and a diversity of somatic cells, including neurons, in juvenile and adult *Nematostella* polyps.

## Results

### Conserved GMP gene orthologs are expressed in the germline and extra-gonadal, basiepithelial cells in Nematostella

Aiming to identify GSCs or ASCs at post-larval stages in *Nematostella*, we performed *in situ* hybridization of the *Nematostella* GMP gene orthologs *piwi1, piwi2, vasa1, vasa2, pl10* and *tudor* on gonadal mesentery regions from adult males and females (Figures 1B-C and S1). We confirmed the previously reported expression of *piwi1*, *piwi2*, *vasa1* and *vasa2* (Praher et al., 2017) and of *pl10* in small and larger oocytes and in spermatogonia, which locate to the periphery of the spermaries (Figures 1B-C’, D-D’’ and S1A-C’’’, E-E’’’). In addition, *piwi1*, *piwi2* and *tudor* were expressed outside of the gonad in basiepithelial cells of the septal filament (Figures 1B-C’ and S1A-A’’, E’; arrowheads). Based on their location and relatively small size, these cells are reminiscent of the previously identified, putative PGCs detected by a *Nematostella* Vasa2 antibody (Chen et al., 2020).

### Colocalization of Piwi1 and Vasa2 proteins in gametes and proliferative, basiepithelial cells of the mesentery

We confirmed recent observations that both monoclonal (Praher et al., 2017) and polyclonal NvVasa2 antibodies (Chen et al., 2020) consistently label developing oocytes, spermatogonia and basiepithelial, extra-gonadal cells in the septal filament and at the distal end of the retractor muscle (Figure S2A-B’’). Using the monoclonal antibody throughout this work, we found that Vasa2+ basiepithelial cells were partly EdU+ after a 3 days-long pulse and concentrated in the reticulate tract (Figure 1H and S3G, H). They were also found as EdU+ duplets in the gonad tract, suggesting that they have recently divided (Figure S3G, I). Vasa2+ cells were not only present in the gonadal region but extended towards pharyngeal and physal regions along the oral-aboral axis of all mesenteries. Within the mesentery, they consistently located to the base of the septal filament and the distal end of the retractor muscle (Figure S3B-E’’’).

To test if Vasa2+ cells coexpress Piwi1 protein, as suggested by *piwi1* mRNA expression (Figure 1B-C’), we generated a CRISPR/Cas9-mediated transgenic knock-in allele (*piwi1^mOr2^)* consisting of the mOrange2 (mOr2) fluorophore fused to the N-terminus of the Piwi1 protein (mOr2-Piwi1; Figure 1D). We found that Vasa2 and mOr2-Piwi1 proteins colocalize to growing oocytes and to EdU+, putative spermatogonia (Figure 1E-G’) (Tucker et al., 2011), confirming that Piwi1 and Vasa2 proteins are coexpressed in developing gametes. In growing oocytes, mOr2-Piwi1 and Vasa2 proteins colocalized together with *tudor* and *piwi1* mRNAs to perinuclear germ granules (“n"; Figures 1G-G’ and S4) (Chen et al., 2020; Praher et al., 2017). In addition, we showed that indeed, basiepithelial Vasa2+ cells coexpress mOr2-Piwi1 protein all along adult and juvenile mesenteries (Figures 1E-E’ and 2, arrowheads). While in adults, some Vasa2+/Piwi1+ cells were labelled after a 3 days-long EdU pulse and are thus proliferative, a larger proportion of Vasa2+/Piwi1+ stayed non-labelled and may thus represent quiescent or slowly cycling cells (Figure 1H-H’, asterisks).

Despite the absence of Vasa2 immunostaining, *mOr2* (Figure S8A) or *piwi1* mRNA signal, we found low levels of mOr2-Piwi1 fusion protein in epidermal cell patches (Figures 2F-F’ and S7A, C), suggesting that epidermal *mOr2-piwi1* mRNA is expressed below our detection limits. We concluded that Piwi1 and Vasa2 proteins are coexpressed in developing gametes and in extra-gonadal, basiepithelial Vasa2+/Piwi1+ cells that locate along specific tracts of the mesenteries in juvenile and adult polyps.

**Figure 2.**
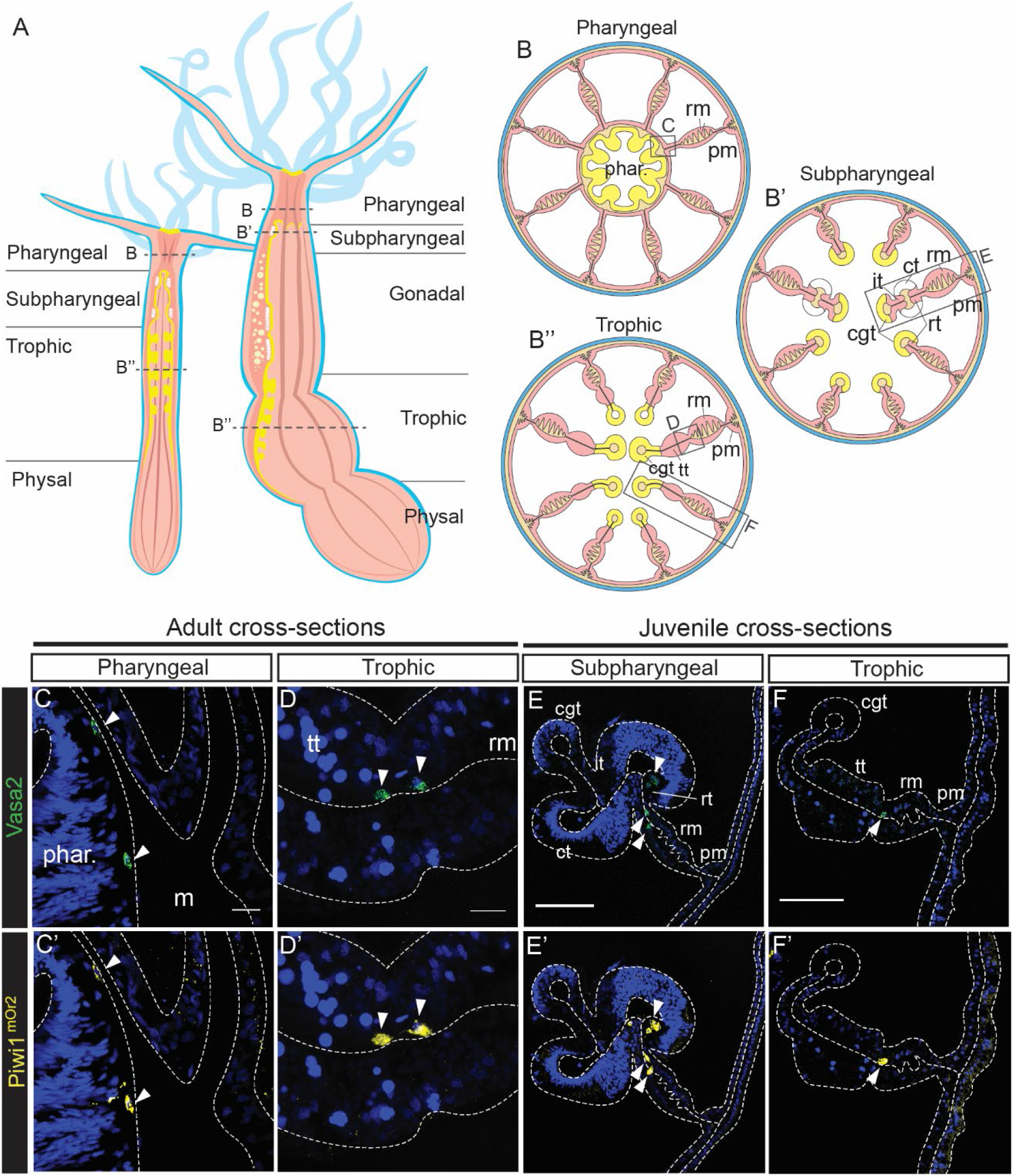
Extra-gonadal location of Vasa2+/Piwi1+ basiepithelial cells in adult and juvenile mesenteries. (A-B’’) Schematics of longitudinal sections of a juvenile and an adult polyp highlighting their mesentery anatomy (A) and cross sections across the pharyngeal (B), subpharyngeal (B’) and trophic (B’’) regions, which have similar morphology in juvenile and adults. (C-F’) Confocal imaging stacks of cross sections of extra-gonadal adult (C-D’) and juvenile (E-F’) mesentery regions immunostained for Vasa2 (C-F, green) and mOr2-Piwi (C’-F’, yellow) as indicated in B-B’’. Vasa2 and mOr2-Piwi1 colocalize to basiepithelial cells of the adult pharynx (C, C’, arrowheads), the adult and juvenile trophic region (D, D’, F, F’, arrowheads), and the juvenile subpharyngeal region of the mesentery (E, E’, arrowheads). Blue: Hoechst DNA dye. phar: pharynx; cgt: cnidoglandular tract; it: intermediate tract; ct: ciliated tract; rt: reticulate tract; tt: trophic tract; rm: retractor muscle; pm: parietal muscle. Scale bars: 10 μm (B, C) and 50 μm (D, E).

### Lineage tracing of Vasa2+/Piwi1+ cells highlights their somatic progeny in adults and juveniles

We studied the potential of extra-gonadal Vasa2+/Piwi1+ cells by generating transgenic reporter lines for *piwi1* and *vasa2* that allowed lineage tracing. First, we characterized an I-SceI-meganuclease-mediated *vasa2*∷mOr2 transgenic line that drives *mOr2* expression from a 1,6kb long upstream region of the *Nematostella vasa2* gene. Codetection of *vasa2*∷mOr2 and Vasa2 proteins in growing oocytes (Figure S5B-B’), spermatogonia (Figure S5C-C’, D) and extra-gonadal cells (Figures S5F-F’) showed that the *vasa2*∷mOr2 line reliably reports endogenous Vasa2 expression in adults. In addition, we found high levels of *vasa2*∷mOr2 protein colocalizing with Vasa2 protein to basiepithelial cells at the distal end of the retractor muscle all along the oral-aboral axis of the juvenile mesenteries (Figures 3A-A’’ and S6B-D’), confirming our observations using the *piwi1^mOr2^* line. Strikingly, we detected lower levels of *vasa2* promoter-driven mOr2, but not Vasa2 protein, in cells scattered throughout the body column gastrodermis (Figure S5E-H’’). Many of those mOr2+/Vasa2– were able to proliferate (EdU+; Figures S5E’-H’’, S6E’-H’’ and S7J-K) and located to similar parts of the mesentery and body wall of adults and juveniles. More specifically, mOr2+/Vasa2– cells were found in the cnidoglandular tract (Figure S6E-E’’) and in the highly proliferative (EdU+) distal end of the ciliated tract (Figures S5E-E’’ and S6F-F’’) (Babonis et al., 2019). mOr2+/Vasa2– cells located also along the retractor muscle (Figures S5G-G’’ and S6G-G’’), parietal muscle (Figures 3B-B’’ and S7H) and adjacent parts of the body wall gastrodermis (Figure S7H), reaching to the tentacle base (Figure S7F, arrows). Notably, these gastrodermal cells often exhibited neuron-like shapes (Figures S5G’ and S7J, K) or filopodia-like extensions as typical for migratory cells (Figures S5H’ and S7H). As for *piwi1^mOr2^*, we observed *vasa2*∷mOr2+ cells in partly EdU+ patches throughout the epidermis despite the absence of detectable Vasa2 protein (Figures 3B’, F’, S6C’, D’, H’ and S7E, G, I).

**Figure 3.**
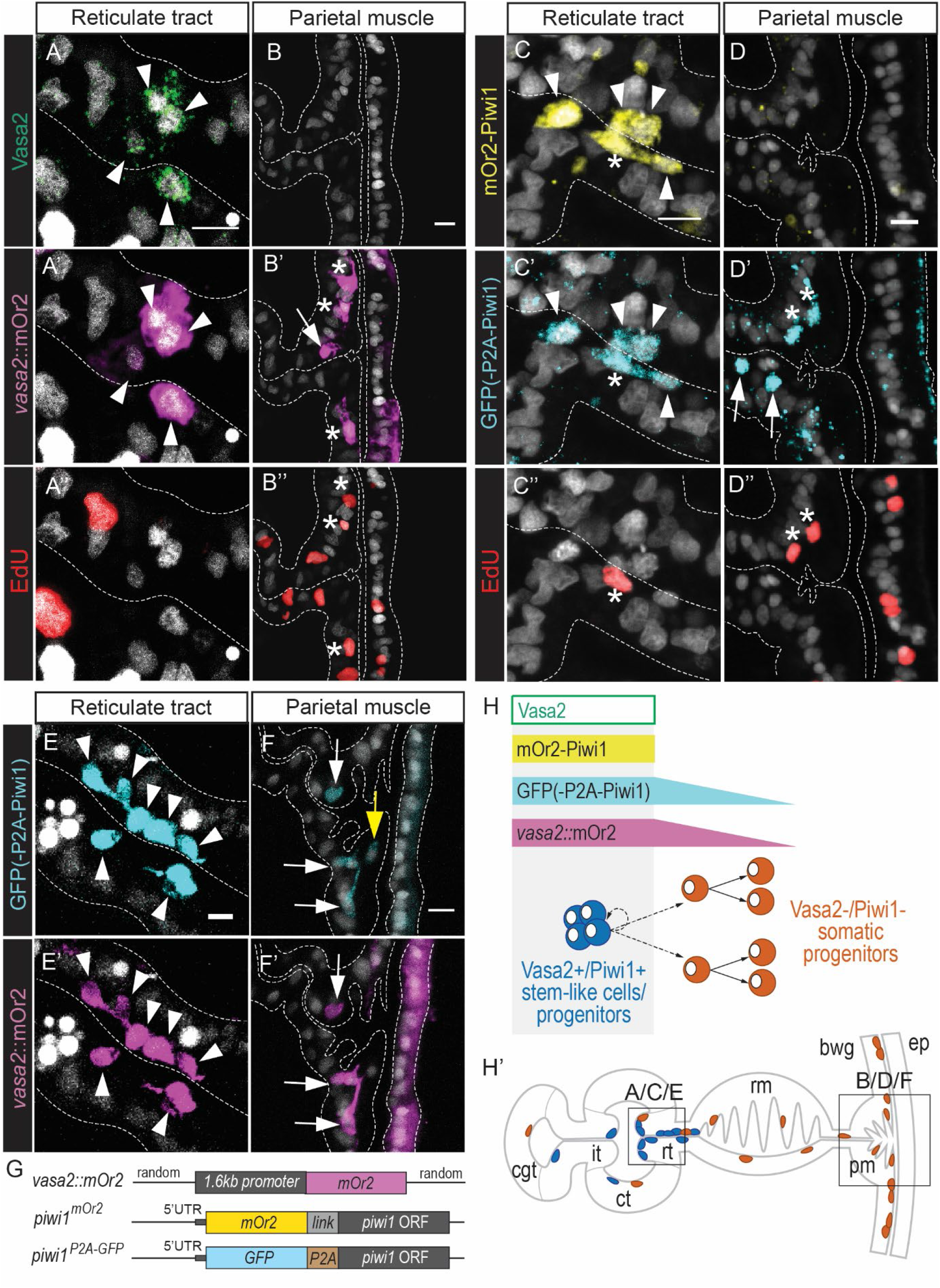
Independent transgenic reporter lines allow tracing somatic, proliferative progeny of Vasa2+/Piwi1+ cells to the parietal muscle tract and body wall gastrodermis. (A-F’) Confocal imaging stacks of cross sections of transgenic juvenile mesenteries immunostained for Vasa2 (A, B, green), mOr2 (A’, B’, E’, F’, yellow), GFP (C’, D’, E, F, cyan) and labelled for S-phase by EdU (A’’, B’’, C’’, D’’, 1 hour pulse, red). Areas of the cross sections shown as in (H’). Vasa2 protein, *vasa2*∷mOr2, mOr2-Piwi1, and GFP(-P2A-Piwi1) colocalize to basiepithelial cells between the septal filament and the retractor muscle of the juvenile subpharyngeal mesentery region (A-A’, C-C’, E-E’, arrowheads), some undergoing S-phase (C-C’’, asterisk). Relatively low levels of *vasa2*∷mOr2 and GFP colocalize to basiepithelial, Vasa2- and mOr2-Piwi1-cells within the parietal muscle tract (F-F’, white arrowheads). Some of these cells undergo S-phase (B-B’’, D-D’’, asterisks). Note that a single GFP+ cell does not colocalize with *vasa2*∷mOr2 in the parietal muscle tract (F, F’, yellow arrowhead). (G) Representation of the reporter line alleles of each transgenic line. (H-H’) Schematics summarizing the location and relative levels of Vasa2 protein and the different reporter fluorophores in potential Vasa2+/Piwi1+ stem cells and their somatic progeny, as well as their location in the subpharyngeal mesentery region (H’). Vasa2 protein, mOr2-Piwi and high levels of *vasa2*∷mOr2 and GFP(-P2A-Piwi1) are found in a population of basiepithelial Vasa2+/Piwi1+ stem-like cells (depicted in dark blue). Lower levels of *vasa2*∷mOr2 and GFP are detected in basiepithelial Vasa2-/Piwi1-cells (depicted in dark orange) likely derived from Vasa2+/Piwi1+ stem-like cells. Grey: Hoechst DNA dye. cgt: cnidoglandular tract; ct: ciliated tract; it: intermediate tract; rt: reticulate tract; rm: retractor muscle; pm: parietal muscle; bwg: body wall gastrodermis; ep: epidermis. Scale bars: 5μm.

As mOr2+/Vasa2– cells could also derive from ectopic expression of the *vasa2* promoter, we tested for ectopic *mOr2* mRNA in *vasa2∷mOr2* reporter animals and generated a CRISPR/Cas9-mediated *piwi1^P2A-GFP^* knock-in allele as an alternative lineage tracing tool. Using *in situ* hybridization, we confirmed that *mOr2* mRNA expression is specific to the germline (Figure S8C, D’, arrows) and to small, basiepithelial cells of the reticulate tract (Figure S8C, D, arrowheads). We generated a *piwi1^P2A-GFP^* knock-in allele that comprises a P2A self-cleaving peptide-encoding sequence (Kim et al., 2011; Tripathi et al., 2022) between the GFP and the N-terminal Piwi1-encoding regions. As this allows cytoplasmic inheritance of GFP to occur independently of Piwi1 protein (Kim et al., 2011; Tripathi et al., 2022), we used *piwi1^P2A-GFP^* to transiently track the progeny of Vasa2+/Piwi1+ in a similar manner to the *vasa2∷mOr2* line. Colocalization of high GFP and mOr2 levels in a *piwi1^P2A-GFP^*/*piwi1^mOr2^* double reporter line further validated the presence of extra-gonadal Vasa2+/Piwi1+ cells along the juvenile mesenteries (Figure 3C-C’’, arrowheads). In addition, we found GFP+/mOr2-Piwi1– cells, often proliferative (EdU+), especially prominent along the parietal muscle region of the gastrodermis (Figures 3D-D’’ and S7B-B’, D-D’), as in *vasa2∷mOr2* animals (Figure S7F, H). These cells thus likely consist of progeny cells from Vasa2+/Piwi1+ mesenterial cells (Figure S7L’). To test if the putative progeny cells overlapped in *piwi1^P2A-GFP^* and *vasa2∷mOr2* lines, we crossed both lines and carefully analyzed their juvenile offspring. As expected, we found high levels of GFP and mOr2 colocalizing in Vasa2+/Piwi1+ cells (Figures 3E-E’ and S9A, B-B’’, arrowheads). Relatively low levels of both reporter proteins also colocalized to cells of the parietal muscle (Figures 3F-F’ and S9B-B’’, arrows) and to both the body wall gastrodermis and epidermis (Figures 3F-F’ and S9F-F’, arrows) of juveniles. This result further supported that both *piwi1^P2A-GFP^* and *vasa2∷mOr2* reporter lines highlight somatic progeny cells derived from the mesenterial Vasa2+/Piwi1+ cells. Additionally, however, we found GFP+/mOr2– cells in the retractor muscle region (Figure S9E, asterisks) and body wall (Figure 3F-F’, yellow arrow; S9F-F’, asterisk). These could be explained by a putative population of Piwi1+/Vasa2– cells that we were unable to detect in *piwi1^mOr2^* animals or by ISH against *piwi1* mRNA. We also found mOr2+/GFP– cells in the cnidoglandular tract (Figure S9C-C’, asterisks) and ciliated tract (Figure S9D-D’, asterisks). mOr2+/GFP– cells may result from relatively higher *vasa2:*:mOr2 levels, partly caused by the integration of *vasa2*∷mOr2 reporter construct concatemers as previously reported for *Nematostella* and fish (Renfer and Technau, 2017; Thermes et al., 2002)

In line with low epidermal levels of mOr2-Piwi1, we also found low, homogeneous levels of GFP throughout the epidermis of *piwi1^P2A-GFP^* animals (Figures S7A’, C’). Interestingly, short, growing tentacle buds showed higher expression of both *piwi1^P2A-GFP^* and *vasa2∷mOr2* reporter levels compared to fully grown tentacles (Figure S7A’, E; arrowheads).

Overall, the large overlap between the cells labelled by *piwi1^P2A-GFP^* and *vasa2∷mOr2* confirms that the progeny of Vasa2+/Piwi1+ cells does not only develop into gametes, but also contributes to a diversity of somatic cells throughout the polyp body column.

### A set of gastrodermal neuron-like cells derives from Vasa2+/Piwi1+ stem-like cells

The parietal muscle region and adjacent body wall gastrodermis in juveniles shows a particularly high concentration of neural-like, putative progeny cells that are double-labelled in *piwi1^P2A-GFP^/vasa2∷mOr2* animals (Figures 3F-F’ and S9F-F’). We therefore tested if Vasa2+/Piwi1+ stem-like cells contribute to neuronal lineages by crossing the *piwi1^P2A-GFP^* line with the *soxB(2)∷mOr2* transgenic line that labels neural progenitor cells in larval and primary polyps (Richards and Rentzsch, 2014). While *soxB(2)-*driven mOr2 protein was absent in Vasa2+/Piwi1+ cells (Figure 4A-A’, arrowheads), we found both *soxB(2):*:mOr2 and GFP coexpressed at low levels in basiepithelial EdU+ cells along the retractor muscle (Figure 4B-B’), parietal muscle (Figure 4D-D’) and in the body wall gastrodermis (Figure 4E-E’). Higher levels of *soxB(2):*:mOr2 were found in GFP– cells displaying shapes of differentiated neurons (Figure 4C-C’, F-F’). Based on these observations, we suggest that a subset of the Vasa2+/Piwi1+ cell progeny develops into SoxB(2)+ neural progenitors and neurons within the retractor muscle, parietal muscle and body wall gastrodermis regions (Figure 4G-G’).

**Figure 4.**
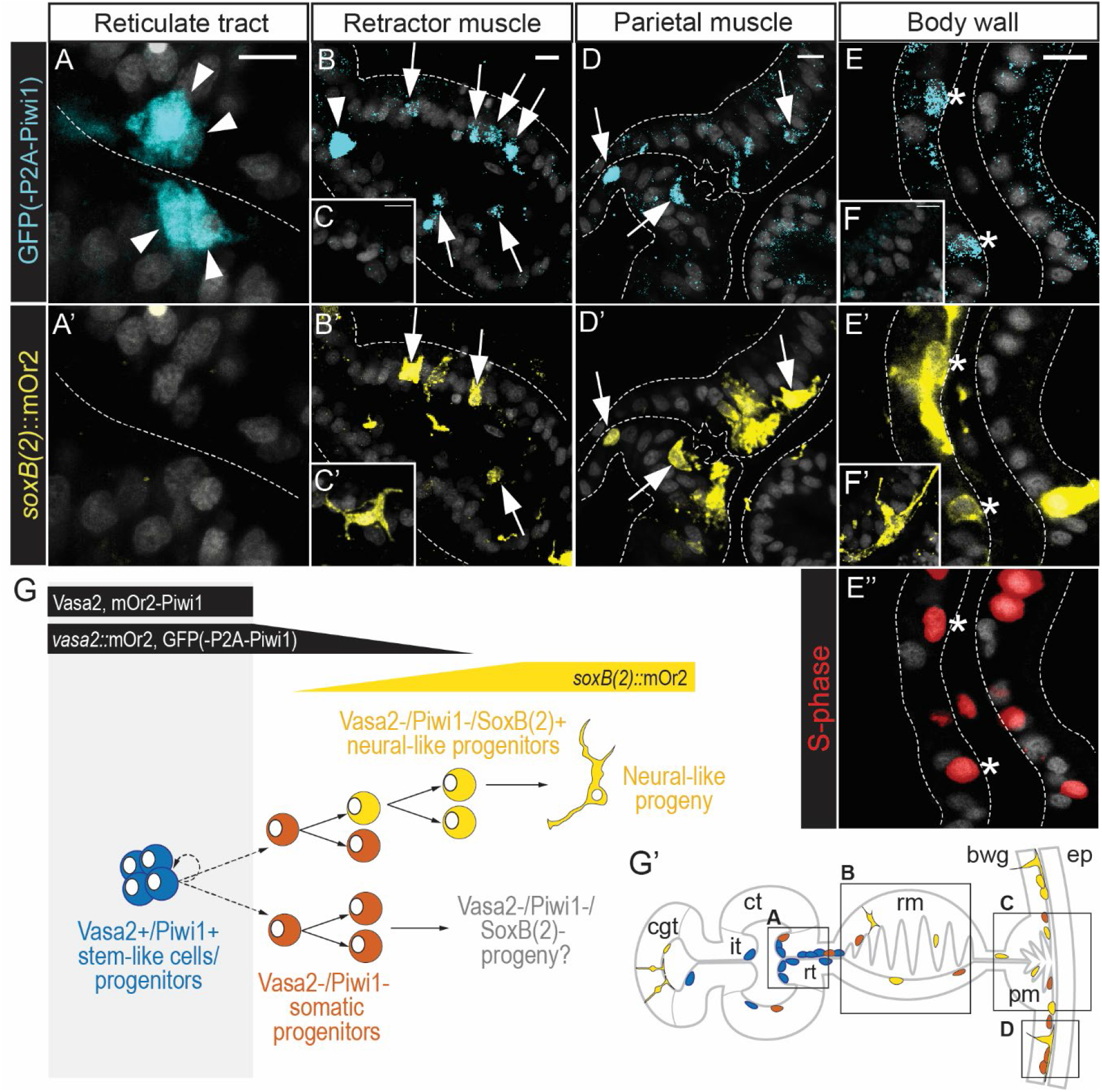
A subset of somatic progeny cells has neuronal fate. (A-F) Confocal imaging stacks of cross sections of juvenile mesenteries from animals carrying both the *soxB(2)*∷mOr2 transgene and a *piwi1^P2A-GFP^* allele. Tissues were immunostained for GFP (A-F, cyan), mOr2 (A’-F’, yellow), and labelled for S-phase by EdU (E’’, 1 hour pulse, red). Areas of the cross sections shown as indicated in G’. Reticulate tract cells with high GFP levels do not express *soxB(2)*∷mOr2 (A, A’, arrowheads), and neurons with high levels of *soxB(2)*∷mOr2 are GFP-(C, C’, F, F’). Colocalization of low levels of GFP and *soxB(2)*∷mOr2 is found in basiepithelial and epithelial cells, some in S-phase (E-E’’, asterisks), in the retractor muscle (B, B’, arrows), parietal muscle (D, D’, arrows) and body wall gastrodermis (E-E’). (G-G’) Schematics depicting the proposed cell lineage (G) and the location within the subpharyngeal mesentery region (G’) of the stem-like cells (dark blue) and their derived *soxB(2)*∷mOr2+ neural progeny (yellow). Potential, alternative *soxB(2)*-fates of the somatic progeny are depicted in dark orange. Indicative cellular levels of Vasa2 protein and *vasa2*∷mOr2, mOr2-Piwi1^mOr2^, and GFP(-P2A-Piwi1) reporters are depicted in black. *soxB(2)*∷mOr2 levels in neural progenitors are depicted in yellow. Grey: Hoechst DNA dye. cgt: cnidoglandular tract; ct: ciliated tract; it: intermediate tract; rt: reticulate tract; rm: retractor muscle; pm: parietal muscle; bwg: body wall gastrodermis; ep: epidermis. Scale bars: 5μm.

## Discussion

Adult PriSCs have been found and molecularly characterized in a few animals that share high body plasticity and regenerative capacities (e.g. planarian neoblasts, hydrozoan i-cells) (Funayama, 2013; Hulett et al., 2022; Varley et al., 2022; Wagner et al., 2011), but their existence and biology is still unknown for many non-bilaterian phyla. Cnidarians are particularly well suited to study the cellular basis and evolution of body plasticity due to their regenerative abilities, phylogenetic position as sister group to bilaterians, and the availability of powerful molecular tools in several cnidarian species (He et al., 2018; Ikmi et al., 2014; Renfer et al., 2010; Sebé-Pedrós et al., 2018; Siebert et al., 2019; Weissbourd et al., 2021; Wittlieb et al., 2006). Currently, however, hydrozoan i-cells are the only known cnidarian adult PriSCs, leaving open if these are ancestral to cnidarians. Identifying and characterizing adult PriSCs in non-hydrozoan cnidarians, such as the sea anemone *Nematostella* vectensis, is therefore informative to understand the evolution of cnidarian and bilaterian stem cell systems, and the cellular biology underlying whole-body regeneration.

In this study, we used Vasa2 immunostainings and a knock-in mOr2-Piwi1 fusion protein allele to identify a population of proliferative cells that coexpress Vasa2 and Piwi1 protein in the sea anemone *Nematostella vectensis*. These cells share a mesenterial location at the basis of the septal filament (i.e., adult reticulate tract) and the distal part of the retractor muscle in both juvenile and adult polyps, corresponding to previously described potential PGCs (Chen et al., 2020). The identification of Vasa2+/Piwi1+ cells in an equivalent place of the mesenteries in *Euphylia ancora* suggests that their location is conserved among hexacorallians (i.e. sea anemones and stony corals).

The presence of Vasa2+/Piwi1+ cells not only in gonadal regions, but all along the mesenteries of juveniles and adults led us to explore if Vasa2+/Piwi1+ cells also contribute to somatic cell types in *Nematostella*. Indeed, by characterizing a promoter-driven *vasa2∷mOr2* and a *piwi1^P2A-GFP^* knock-in transgenic reporter lines, we have shown that Vasa2+/Piwi1+ cells generate both gametes and a diversity of somatic cell types in juvenile and adult polyps (Figure 5). Somatic progeny cells in the gastrodermis are to a large extent proliferative, basiepithelial and display polarized shapes. We thus propose that at least a subset of the progeny derived from Vasa2+/Piwi1+ cells consist of migrating, transit-amplifying progenitor cells. Some of the progeny cells along the muscle tracts and body wall gastrodermis displayed neural-like shapes, suggesting that they develop into neurons. We confirmed this assumption by showing that a subset of progeny cells from Vasa2+/Piwi1+ cells consists of *soxB(2)*-expressing neural progenitor cells. Previous studies using transgenic reporter lines and single cell RNA-seq analysis have shown that multipotent *soxB(2)*+ neural progenitor cells generate gland cells, neurons and cnidocytes during larval and potentially post-larval development (Richards and Rentzsch, 2014; Steger et al., 2022; Tournière et al., 2022). However, the stem cell populations upstream of such progenitors remain so far unidentified. Here, we propose that Vasa2+/Piwi1+ stem-like cells contribute to at least a subset of *soxB(2)*+ neural progenitor cells and neural lineages in the juvenile and adult gastrodermis. Whether Vasa2+/Piwi1+ stem-like cells are also upstream of *soxB(2)*+ cell lineages during larval development remains to be studied.

**Figure 5.**
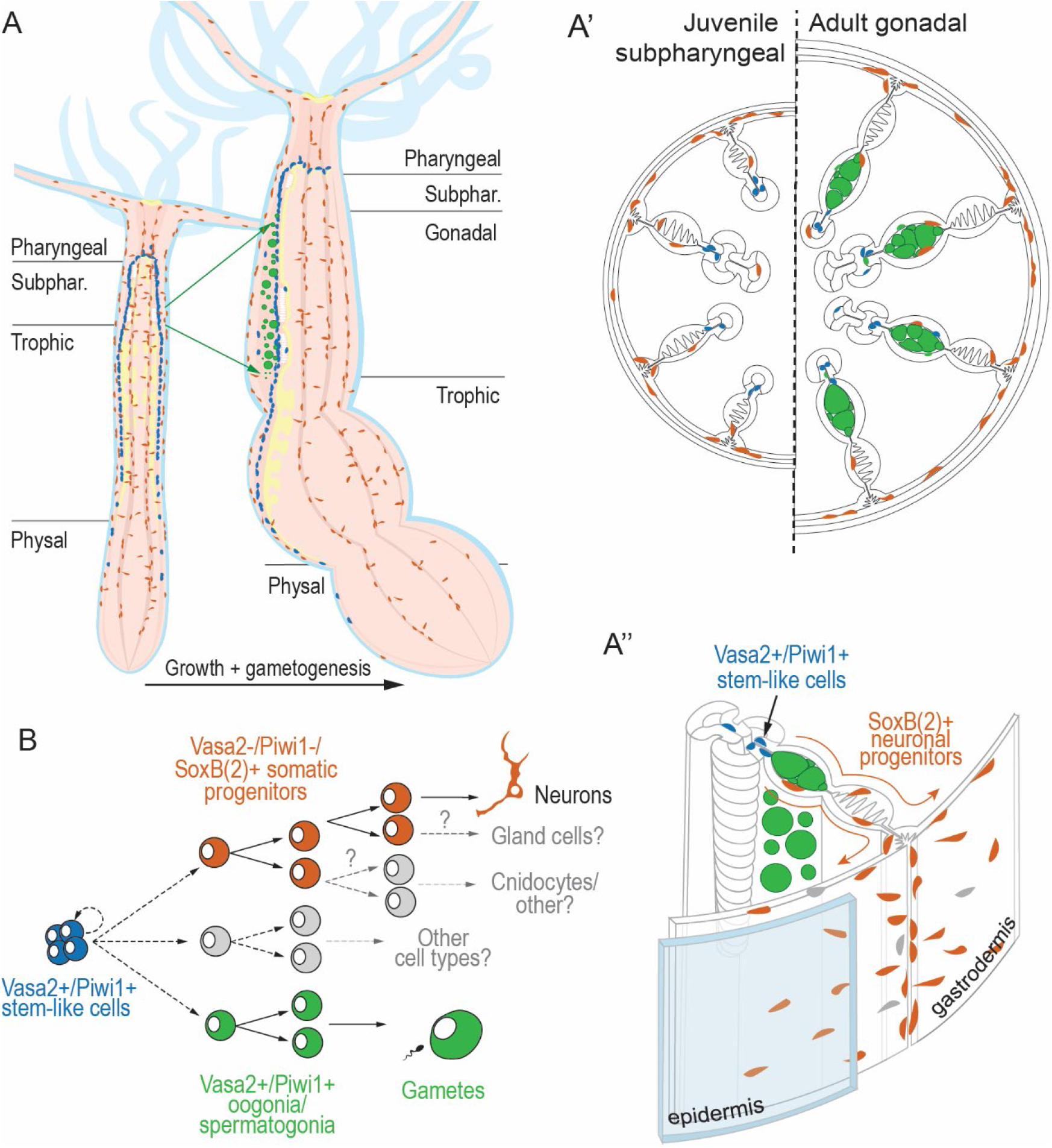
Vasa2+/Piwi1+ stem-like cells in the mesenteries of *Nematostella* as a potential origin for the germline and somatic cell types. (A-A’’) Schematic summary of the results and conclusion, depicting multipotent Vasa2+/Piwi1+ stem-like cells (blue) and their putative somatic/neuronal (orange) and germinal progeny (green) in a longitudinal cross section of a juvenile and an adult polyp (A), cross sections of the juvenile subpharyngeal and adult gonadal mesentery regions (A’) and 3D reconstruction of an adult mesentery including body column (A’’). (B) Proposed cell lineage relationship between an adult Vasa2+/Piwi1+ multipotent stem-like cell population and their germinal and somatic/neuronal progeny.

Our results thus indicate that the mesenterial Vasa2+/Piwi1+ cell population does not consist of PGCs, as previously assumed (Chen et al., 2020; Extavour et al., 2005), but of PriSCs with both germline and somatic potentials. This is in line with recent genomic data that supports a continuous formation of both germline and somatic lineages from a common pool of ASCs through adulthood in stony corals, which are close relatives of sea anemones (López-Nandam et al., 2023; Vasquez Kuntz et al., 2020). Currently, however, the heterogeneity of the *Nematostella* Vasa2+/Piwi1+ cell population remains to be addressed.

While the epidermis shows low levels of GFP or mOr2 proteins in all three independently generated transgenic lines, we have so far failed to detect epidermal *piwi1* or *vasa2* transcript or Vasa2 protein expression. *vasa2* or *piwi1* have also not been identified as markers for epidermal cell clusters of recent single-cell RNA-Seq datasets (Sebé-Pedrós et al., 2018; Steger et al., 2022).These observations are in line with a working hypothesis that epidermal cells express low levels of *vasa2* and *piwi1* and may thus consist of an independent, self-renewing population, similarly to the epidermis of *Hydra* (Hemmrich et al., 2012; Siebert et al., 2019).

While i-cells have so far been considered a hydrozoan-specific trait, the Vasa2+/Piwi1+ stem-like cells described here show strong similarities to hydrozoan i-cells. Both exhibit a relatively small cell size, a high nucleus to cytoplasm ratio, share basiepithelial location (Frank et al., 2009) and express the conserved GMP genes *vasa* and *piwi* (Leclère et al., 2012; Lim et al., 2014; Mochizuki et al., 2001; Rebscher et al., 2008). In addition, both populations generate the germline and neural cell types (David, 2012; Müller et al., 2004; Siebert et al., 2019; Varley et al., 2022). I-cells do not only generate neurons, but also gland cells and cnidocytes (Chrysostomou et al., 2022; Siebert et al., 2019). We have shown that the Vasa2+/Piwi1+ cell population contributes to at least some *soxB(2)*+ cells and neurons in *Nematostella* juveniles. As the reporter fluorophores in our transgenic lines get diluted and degraded in the dividing and differentiating progeny of Vasa2+/Piwi1+ cells, we could not fully reconstruct all their derived cell lineages. Thus, it remains to be addressed if the adult Vasa2+/Piwi1+ stem-like cell population is upstream of all or only a subset of *soxB(2)* lineages in juvenile and adult *Nematostella* polyps, and whether these derive cnidocytes and gland cells at these stages as i-cells do.

ASCs holding germinal and somatic potential (i.e. adult PriSCs) have been consistently found in animals displaying high body plasticity and regenerative abilities, such as sponges (archeocytes and choanocytes), hydrozoans (i-cells), acoels and planarians (neoblasts), and colonial ascidians (hemoblasts) (Brown et al., 2009; De Mulder et al., 2009; Fierro-Constaín et al., 2017; Frank et al., 2009; Nishimiya-Fujisawa and Kobayashi, 2012; Srivastava, 2022; Sunanaga et al., 2010). Yet, their dispersed phylogenetic occurrence, and the lack of molecular data from most non-bilaterian phyla hampers our understanding of adult PriSCs evolution. In this study, we identify putative adult PriSCs for the first time in a non-hydrozoan cnidarian. Given their partly shared genetic signatures and potentials, we propose that *Nematostella* Vasa2+/Piwi1+ adult stem-like cells are homologous to hydrozoan i-cells. As high body plasticity and regenerative abilities are common across cnidarians, we predict that *vasa+/piwi*+ adult PriSCs are present beyond anthozoans and hydrozoans. The discovery of potential adult PriSCs in *Nematostella* sets the basis for uncovering the cellular and molecular mechanisms that underlie body plasticity in sea anemones and corals, and for further elucidating the evolution of animal stem cell systems.

## Materials and methods

### Animal culture

*Nematostella vectensis* polyps derive from CH6 females and CH2 males constituting the original culture (Hand and Uhlinger, 1992). Female and male adult animals are maintained in separate boxes in the dark at 18°C in 1/3 filtered sea water (*Nematostella* medium, NM), daily fed with fresh *Artemia nauplii*. Spawning is induced every three weeks by a light and temperature shift (25 °C) overnight (12 hours), as previously described (Fritzenwanker and Technau, 2002). Embryos are raised at 21°C, and daily feeding with *Artemia nauplii* is incorporated 7 days post-fertilization, approximately.

### Meganuclease-mediated generation of a vasa2 reporter line

A piece of promoter comprising 1.6kb upstream of *vasa2* 5’ UTR was selected takinginto account the genome wide histone marks H3K27ac, H3K4me1, H3K4me2, H3K4me3 and ChIP-Seq data for P300 (transcriptional coactivator) in adult females (Schwaiger et al., 2014). The promoter fragment was successfully cloned into an existing pJet1.2 plasmid backbone (ThermoFisher) containing an mOrange2 (mOr2) ORF sequence and SceI sites. The resulting vector consisted of 1.6kb *vasa2* promoter upstream of mOr2, altogether flanked by inverted I-SceI sites for I-SceI meganuclease-mediated integration into the genome. Microinjection was performed as previously described (Renfer et al., 2010). 150 injected zygotes were raised until juvenile stage, then selected if presenting gastrodermal mOr2 signal. At adult stage one single animal displaying mOr2 in its developing gametes was selected as a founder and outcrossed with wild type to stablish the F1 generation.

### CRISPR-Cas9 mediated generation of piwi1 knock-in lines

A previously published protocol for the generation of CRISPR/Cas9-mediated knock-in lines in *Nematostella* (Lebouvier et al., 2022) was used and adapted as follows. Two guide RNA (gRNA) target regions with putative cutting sites located 174bp upstream or 229bp downstream of the Start codon of the *Nematostella piwi1* (v1g79423) were designed using CRISPOR (Concordet and Haeussler, 2018). Templates for gRNAs were generated using annealed and PCR-amplified oligos (Bassett et al. 2013): two T7-and gRNA-encoding oligos (ThermoFisher, desalted): 5’-GAAATTAATACGACTCACTATAGacaccacggttccaatatggGTTTTAGAGCTAGAAATAGCAAG-3’ (upstream of Stop codon); 5’-GAAATTAATACGACTCACTATAGGATGTCAGTGGAACCACGTGGTTTTAGAGCTAGAAATAGCAAG-3’ (downstream of Stop codon); invariant reverse primer (ThermoFisher, desalted): 5’-AAAAGCACCGACTCGGTGCCACTTTTTCAAGTTGATAACGGACTAGCCTTATTTTAACTTGCTATTTCTAGCTCTAAAAC-3’.

Guide RNAs were *in vitro* transcribed using a T7 MegaScript transcription kit (ThermoFisher) followed by ammonium chloride precipitation and diluted in nuclease-free H2O to a final concentration of 1,5μg/μl. The DNA donor fragment for homology-mediated repair (HDR) consisted of the coding sequences for the mOr2 protein followed by a GGGGS^2^ linker (mOr2-Piwi1) or a GFP protein followed by a P2A sequence (GFP-P2A-Piwi1) cloned in frame to the 5’ end of the *piwi1* gene open-reading frame. In addition, the construct included a 995bp long ‘left’ homology arm (upstream of Piwi1 coding region) and a 1184bp ‘right’ homology arm (downstream and overlapping the Piwi1 coding region). The entire construct was cloned into a pJet1.2 plasmid backbone (ThermoFisher) using Gibson assembly master mix (NEB) (Gibson et al., 2009). Donor DNA fragment was PCR-amplified using 5’-biotin-labelled oligos flanking the donor fragment to increase integration efficiency as previously described for medaka fish (Gutierrez-Triana et al., 2018). Oligo sequences (ThermoFisher; desalted) are: Forward oligo 5’-[Biotin]-TACGACTCACTATAGGGAGAGCGGC-3’; reverse oligo 5’-CCATGGCAGCTGAGAATATTGTAGGA-3’.

The Cas9-mediated knock-in injection was performed by modifying previously described protocols for CRISPR/Cas9-mediated mutagenesis and used 0,75μg/μl nls-Cas9 protein (PacBio), 75ng/μl of each guide RNA, 70ng/μl column-purified donor DNA, and modified injection buffer containing 220mM KCl (Burger et al., 2016; Kraus et al., 2016). After injection of approx. 1600 zygotes and subsequent raising, juvenile polyps displaying mOr2 or GFP signal in their gastrodermis were selected and raised until sexual maturity. At the adult stage, polyps presenting mOr2 or GFP signal in the developing gametes were selected as founder animals and outcrossed with wild type animals to generate F1 generation animals. The successful integration of the GFP-P2A-Piwi1 fragment knock-in was validated on the transcript level by *in vitro* synthesis of cDNA based on extracted total RNA from a mix of 3 days old larvae resulting from a cross of two heterozygous F1 animals. Based on this cDNA, the transitions between the 5’ UTR and GFP-encoding region, and between the P2A peptide-encoding region and the 5’ end of the Piwi1 open-reading frame have been amplified using oligos outside of the ‘homology arm’ regions, if possible (exception: short Piwi1 5’UTR was fully included within homology arm). Sanger sequencing of gel-eluted and column-purified PCR fragments confirmed the flawless integration of the GFP-P2A fragment directly upstream and in frame with the Piwi1 open-reading frame. Oligos used for PCR & sequencing of GFP-P2A-Piwi1: Piwi1-T-Forward-1: 5’-GAGAGACAGAGCGTTTGAGAGAGAGAC-3’; Piwi1-T-Forward-2: 5’-GAGTTGTGTAATTTTAGGTAAGTTTT GGTC-3’; Piwi1-T-Reverse-1: 5’-TTCCTTCCCACTTGCTTCATGTC-3’; Piwi1-T-Reverse-2: 5’-GTAACCTGGCCACAGCTCAAGC-3’; GFP-Forward (used with Piwi1-Reverse-1 or -2): 5’-TCCGCCCTGAGCAAAGACC-3’; GFP-Reverse (used with Piwi1-Forward-1 or -2): 5’-TTGCCGTAGGTGGCATCGC-3’. The successful and flawless integration of the mOr2-Piwi1 knock-in was validated on the genomic level by extracting genomic DNA from mixed tentacle clips of several homozygous F2 animals from the same founder F1 animals. Based on this gDNA, the transitions between the 5’ UTR of the *piwi1* gene and mOr2, and between the GGGGS^2^ linker and 5’ end of the Piwi1 ORF have been amplified using oligos outside of the ‘homology arm’ regions. Gel elution and Sanger sequencing confirmed the flawless integration. Oligos used: Piwi1-G-Forward: CGCGATACACACTAAACATCTAGGC; Piwi1-G-Reverse: GCAAAAGATAGGACAATCAGCCCCATC; mOrange2-Reverse (used with Piwi1-G-Forward): CTGTCTGAAAGCCCTCGTATGGTC; mOrange2-Forward (used with Piwi1-G-Reverse): CAAGGCAAAGAAGCCAGTGCAGC.

### Gene cloning and RNA probe synthesis

A set of *Nematostella* GMP gene orthologs was chosen from (Fierro-Constaín et al., 2017), comprising *piwi1*, *piwi2*, *vasa1*, *vasa2*, *pl10* and *tudor*. RNA of whole adult female *Nematostella* was extracted using Trizol (ThermoFisher) as indicated by the manufacturer. cDNA was synthetized using the SuperScript III First-Strand Synthesis System (Thermo Fisher), followed by PCR amplification of fragments of the genes of interest. Primers designed with Primer3Plus (http://www.bioinformatics.nl/cgi-bin/primer3plus/primer3plus.cgi). Cloned gene fragments were inserted into the pGEM-T Easy vector (Promega, A1360) and transformed into One Shot Top 10 chemically competent *E. coli* (Invitrogen). Sanger sequencing verified the cloned sequences at the sequencing facility of the Department of Biological Sciences, University of Bergen, Bergen, Norway. Digoxygenin (DIG)-labelled antisense riboprobes were generated using a T7 or SP6 MEGAscript Kit (Invitrogen, AMB1334/AMB1330) and DIG RNA Labelling Mix (Roche) according to the manufacturers’ protocols and (Genikhovich and Technau, 2009). Oligos used for PCR: *piwi1* and *piwi2* as in (Praher et al., 2017); *vasa2* and *tudor* as in (Chen et al., 2020); *vasa1*-Forward: CCCAAACCAAGCCAACCAAGGC, *vasa1*-Reverse: ATGTCAAGGC CACGAGCAGC; *pl10*-Forward: CTTAGCAGGATCTACATGGAAGGGC; *pl10*-Reverse: CCAGTCCTGACCACCGCTG; mOr2-Forward: ATGGTGAGCAAGGGCGA GG; mOr2-Reverse: GTTCCACGATGGTGTAGTCCTCG.

### Colorimetric *in situ* hybridization

Adult female polyps kept in NM were left to relax in petri dishes, then MgCl2 was carefully added and mixed in until reaching a 0.1M MgCl2/NM solution. Animals were left to relax for 30 minutes. The body cavity of the animals was flushed with this medium through the mouth opening to ensure full extension of the body column and mesenteries. The polyps were then transferred to a new dish containing 3.7% Formaldehyde NM solution, and head and physa were cut off with a scalpel. The body column was opened longitudinally with microdissection scissors to ensure penetrance of the fixative. The tissue was gently transferred into 1xPBS/3.7% Formaldehyde/0.5% DMSO/0.1% Tween20 to fix overnight at 4°C. Afterwards, mesenteries with or without body wall were dissected in fixative and cut in approx. 3-5mm long pieces. The tissue pieces were washed in 1x PBS/0.1% Tween20, followed by several washes in 100% methanol until the brown pigment was completely removed. Tissue pieces were finally stored in 100% methanol at −20°C.

The urea-based *in situ* hybridization protocol was adapted from (Sinigaglia et al., 2018), with a series of changes as previously described (Lebouvier et al., 2022). After progressive rehydration in MeOH/PTx (0.3% Triton X-100 in 1xPBS pH 7.4), the tissue was digested with Proteinase K 2.5μg/ml for 5 minutes at room temperature. An additional fixation step (2 min in 0.2% glutaraldehyde/PTx followed by 1 hour in 3.7% formaldehyde/PTx) was added before overnight blocking in a hybridization mix consisting of 50% 8M urea, 5x SSC pH 4.5, 0.3% Triton X-100, 1% SDS, 100 μg/ml heparin and 5 mg/ml Torula yeast RNA. Background staining could be reduced by adding 5% dextrane sulfate (MW > 500,000, Sigma-Aldrich) and 3% Blocking Reagent (Roche) to the hybridization mix during overnight blocking and hybridization of the probe, and hybridization occurred over two days. The probe concentration was 0.75 ng/μl. Stringent washes varied between 2x SSC/0.3% Triton X-100 (SSCT) and 0.1x SSCT. After the SCCT washes, the tissue pieces were washed with 1x PBS/0.1%BSA/0.3% Triton X-100. Pieces of tissue were incubated in 500 ml of NBT/BCIP solution (4.5 μl/ml NBT, 3.5 μl/ml BCIP in Alkaline Phosphatase buffer) (Roche 11383213001 and 11383221001) in the dark from some minutes to several hours, depending on the signal. Once stained, the pieces of tissue were quickly washed in 100% ethanol for 3 minutes, then twice in PBTx and finally stored in 80% glycerol at 4°C.

Before proceeding with the vibratome sectioning overview pictures of the stained pieces of tissue embedded in glycerol were taken in the microscope Leica M165 FC with the Leica DFC450 C camera using the Leica Application Suite X (LAS X) software.

### Immunofluorescence and S-phase labelling

Wild-type and F1 transgenic reporter animals at adult and juvenile (45 days after fertilization, approx.) stages were selected and fixed 24 hours after the last feeding event. To label cells in S-phase, animals were placed in freshly prepared 100μM EdU (Invitrogen) in 2%DMSO NM before fixation. Incubation time was one hour for juveniles, and three days for adults (medium was replaced daily). Animals were relaxed using MgCl2, then fixed in 3.7% Formaldehyde NM for one hour at room temperature and dissected in this same solution in a petri dish. Fixative was washed thoroughly in 1x PBS/0.2% Tween20, followed by a series of dehydration washes (20-50-100% methanol in 1x PBS/0.2% Tween20. 100% methanol washes were applied until pigment was completely removed from the tissue. Samples were stored in 100% methanol at −20°C. After progressive rehydration of the tissue in 1xPBS/0.2% Triton X-100, for samples incubated in EdU a click-it reaction was performed using the reagents of a Click-iT™ EdU Cell Proliferation Kit for Imaging (Invitrogen), following the manufacturer’s protocol. After a 30 minutes incubation in the click-it staining reaction, pieces were washed in in 1xPBS/0.2% Triton X-100.

For the immunofluorescence, tissue pieces were blocked in 1xPBS/10%DMSO/ 5%NGS/0.2% Triton X-100 for two hours at room temperature. Primary antibody incubation was performed in 0.1%DMSO/5%NGS/0.2% Triton X-100 overnight at 4°C using the following antibodies: rabbit anti-Vasa2 1:1000 (Praher et al., 2017), mouse anti-Vasa2 1:500 (Chen et al., 2020), rabbit anti-DsRed 1:100 (Takara Bio Clontech 632496), mouse anti-mCherry 1:100 (Takara Bio Clontech 632543), mouse anti-GFP 1:250 (Abcam Ab1218), rat anti-tubulin (YL1/2) 1:100 (Abcam Ab6160). After washes in 1xPBS/0.2% Triton X-100, tissue was blocked in 1xPBS/5%NGS/0.2% Triton X-100 for 30 minutes at room temperature. Hoechst or DAPI nuclear staining (ThermoFisher) and secondary antibody incubation was performed in 1xPBS/5%NGS/0.2% Triton X-100 overnight at 4°C using the following antibodies: goat-anti-mouse-Alexa488/568 (LifeTech A11001, A11004), goat-anti-rabbit-DyLight488 (LifeTech 35552), goat-anti-rabbit-Alexa568/647 (LifeTech A11011, A21244) and goat-anti-rat-Alexa633 (LifeTech A21094). Finally, tissue pieces were washed thoroughly in 1xPBS/0.2% Triton X-100 and stored at 4°C until sectioning.

### Vibratome sectioning

Vibratome sectioning was performed as previously published (Helm and Capa, 2015), with a series of changes. Stained pieces of mesenteries were selected for embedding in gelatin-albumin medium prepared in advance (22.5 ml PBS (10x), 1.1 g of gelatin Type A Sigma G1890, 67.5 g of albumin Sigma A3912 and 225 ml H20). Pieces were transferred into a plastic mold containing a freshly prepared mixture of 1/4 37%FA gelatin-albumin medium and let solidifying O/N at 4°C in a sealed humid chamber. The solidified gelatin blocks were removed from the mold and placed in PTw to avoid drying. Sectioning was performed with the vibratome Leica VT1000 S, speed = 4, frequency = 6, and section size 20 μm. Slices were stored in PTw in a well plate at 4°C until subsequent mounting and imaging.

### Transmitted light and confocal imaging

*In situ* hybridization and immunofluorescence gelatin slices were mounted on slides (Electron Microscopy Sciences 63418-11) with glycerol and sealed with coverslip (Menzel-Gläser 18×18mm) and clear nail polish. Transmitted light pictures of ISH cross sections were taken in the microscope Nikon Eclipse E800 (60x oil) with the Nikon Digital Sight DS-U3 camera, NIS-Elements software. Immunofluorescence whole mount tissue pieces and cross sections were imaged either on a Leica SP5 confocal microscope (standard PMT detectors, 20x/40x/63x oil-immersion objectives), or an Olympus FV3000 confocal (standard PMT detectors, 40x/63x silicon-immersion objectives). Live juvenile polyps relaxed in MgCl2 were mounted and imaged on the Olympus FV3000 confocal. Transmitted light images and confocal stacks were processed, cropped and adjusted for levels and color balance with Fiji.

## Author contributions

P.M.-P. and P.R.H.S. designed the study and interpreted the results. P.M-P. performed ISH, EdU labelling and immunostainings. P.R.H.S. generated the CRISPR-Cas9 knock-in *piwi1^mOr2^* and *piwi1^P2A-GFP^* constructs and the *piwi1^P2A-GFP^* line. P.M-P generated the *vasa2∷mOr2* construct and line, and the *piwi1^mOr^* line. Transgenic reporter lines were raised, established, and characterized by P.M-P. P.M-P. and P.R.H.S. wrote the manuscript together.

## Acknowledgements

We thank U. Technau’s lab for sharing a monoclonal rabbit anti-NvVasa2 (Praher et al. 2017) and M. Gibson’s lab for sharing a polyclonal rabbit anti-NvVasa2 (Chen et al. 2020). We thank Marie Montjouridès for technical support, which contributed to the characterization of the *vasa2∷mOr2* and *piwi1^mOr2^* reporter lines. We thank P. Steinmetz’s lab members for helpful discussions. We thank F. Rentzsch’s lab for providing the *soxB(2)∷mOr2* reporter line and for constructive feedback.

## SUPPLEMENTARY FIGURES

**Figure S1.**
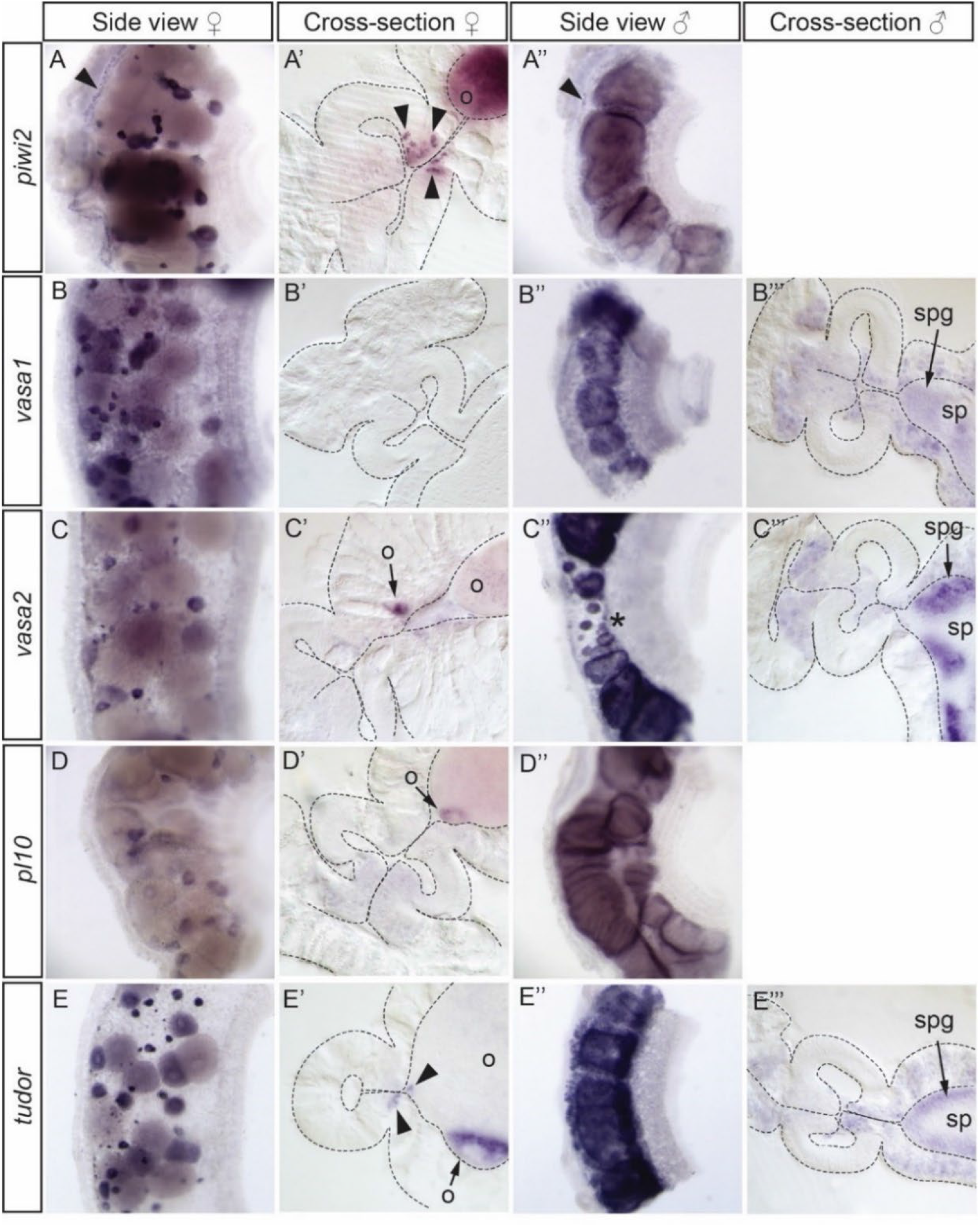
Expression of *Nematostella* GMP marker gene orthologs in the adult gonadal mesentery. Whole mount tissue pieces and cross sections of female and male gonadal mesentery regions stained by ISH for *piwi2* (A-A’’), *vasa1* (B-B’’’), *vasa2* (C-C’’’), *pl10* (D-D’’) and *tudor* (E-E’’’) genes. Expression is found in oocytes of different sizes (o) and in spermatogonia (spg). In males, certain regions of the gonad present smaller spermaries (C’’, asterisk). Expression of *piwi2* and *tudor* is also detected in basiepithelial cells that align and concentrate along the reticulate tract of the mesentery (A-A’’ and E’, arrowheads). o: oocyte; spg: spermatogonia; sp: sperm.

**Figure S2.**
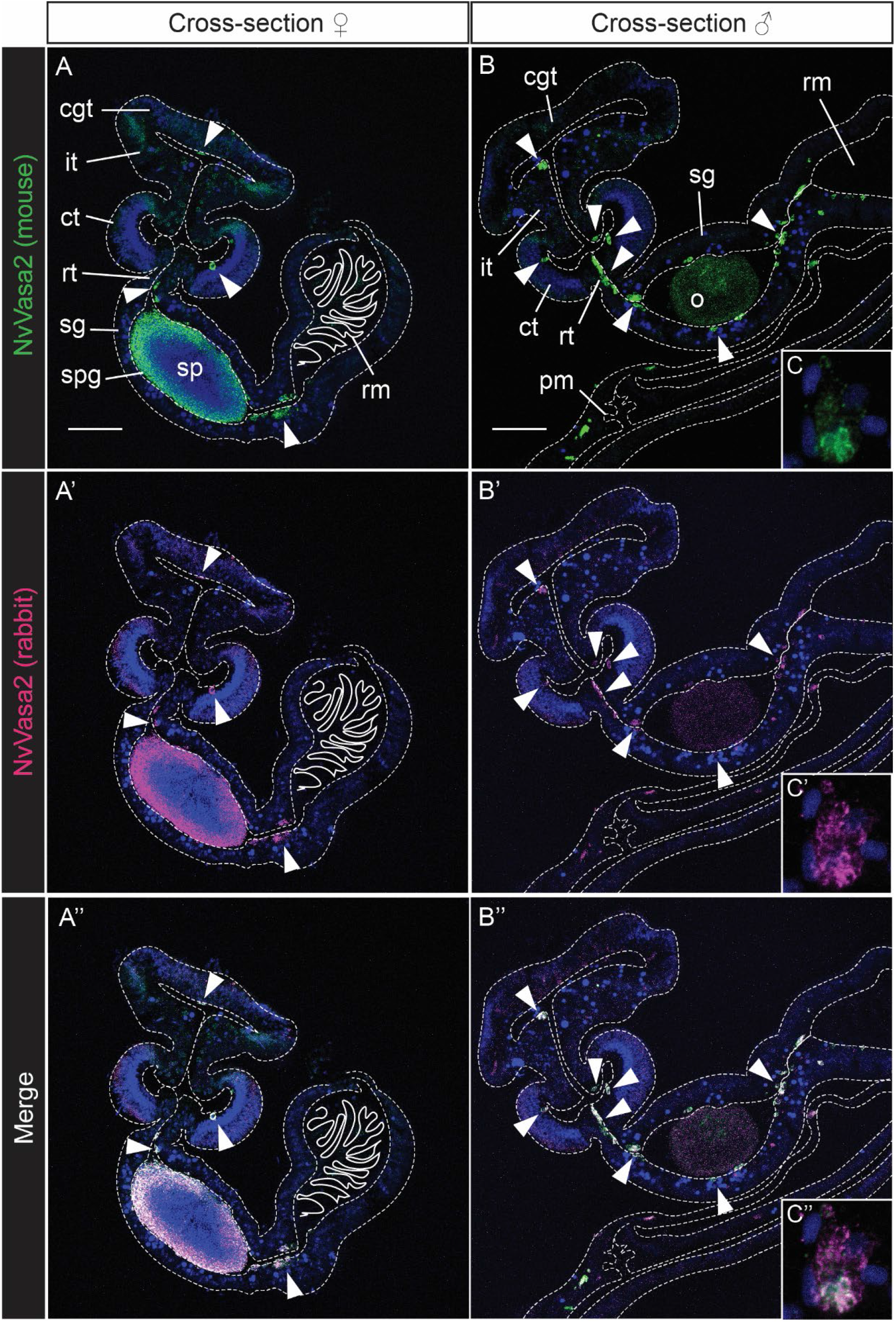
Colocalization of two independently generated antibodies raised against *Nematostella* Vasa2. (A-C’’) Confocal imaging stacks of cross sectioned adult male (A-A’’) and female (B-B’’) gonadal mesentery regions immunostained for Vasa2 with a monoclonal mouse anti-NvVasa2 (green, A, B, C) (Praher et al., 2017) and a monoclonal rabbit anti-NvVasa2 (magenta, A’, B’, C’) (Chen et al., 2020). Complete colocalization is found in developing spermatogonia (spg), oocytes (o) and in basiepithelial cells (arrowheads) located in the septal filament, gonad tract, and in the transition zone between gonad and retractor muscle (rm) (A-B’’, arrowheads). (C-C’’; inlets of B-B’’) Example of unspecific binding of both antibodies to vesicles of a putative mucus cell in the parietal muscle tract. Blue: Hoechst DNA dye. cgt: cnidoglandular tract; it: intermediate tract; ct: ciliated tract; rt: reticulate tract; sg: somatic gonad; o: oocyte; spg: spermatogonia; sp: sperm; sg: somatic gonad, rm: retractor muscle, pm: parietal muscle. Scale bars: 50μm.

**Figure S3.**
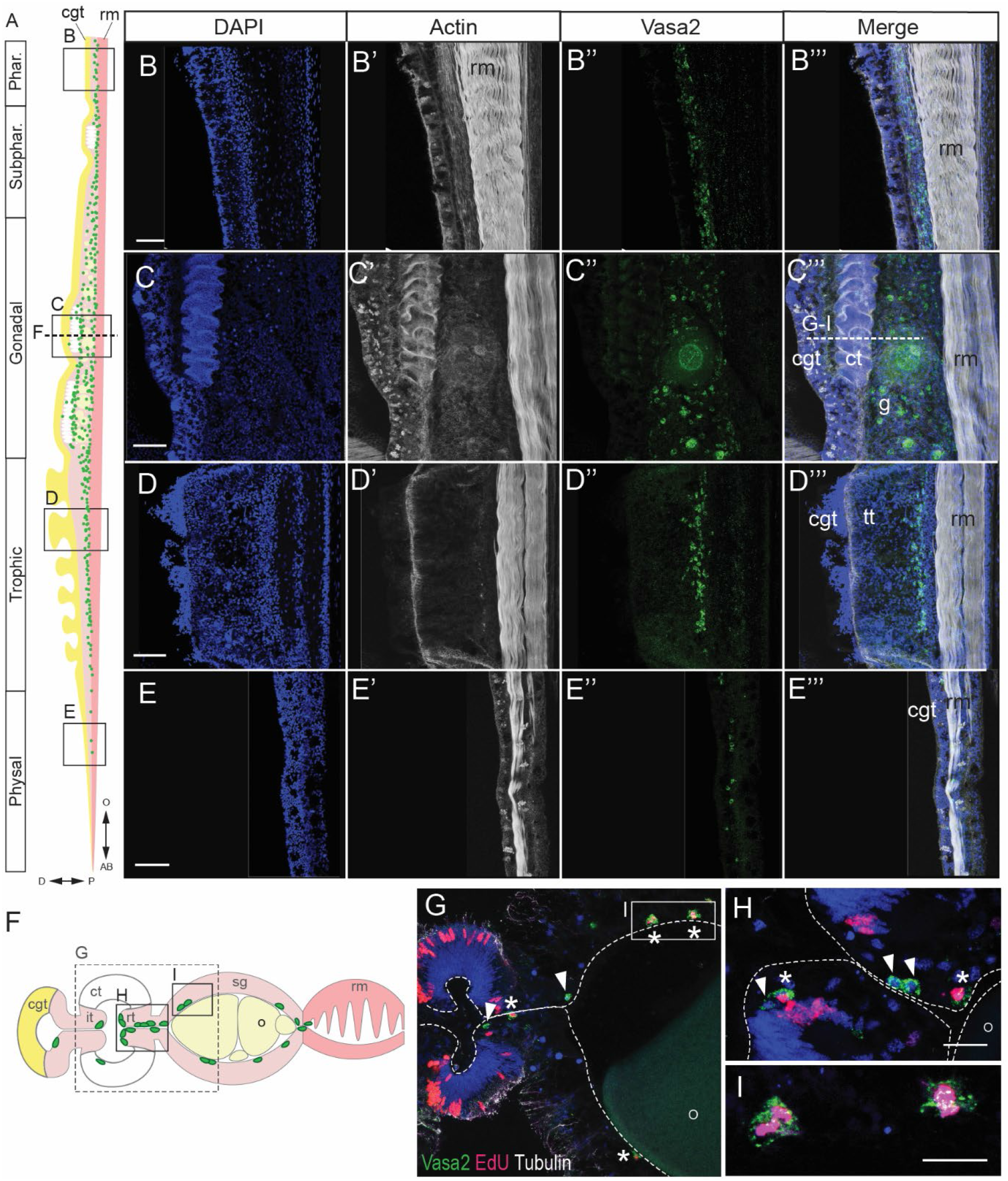
Dividing, basiepithelial Vasa2+ cells populate the mesenteries along the oral-aboral axis. (A, F) Schematics highlighting distribution of Vasa2+ cells (green dots) along oral (O) to aboral (AB) regions (A, side view) or at a cross-section of the gonadal region (F) in an adult female mesentery. Vasa2+ cells locate between the cnidoglandular tract (yellow, ‘cgt’) and the retractor muscle (dark pink, ‘rm’). (B-E’’’) Confocal imaging stacks of pharyngeal (B-B’’’), gonadal (C-C’’’), trophic (D-D’’’) or physal (E-E’’’) regions of a whole adult female mesentery immunostained for F-actin (white) and Vasa2 (green). Note the dense concentration of F-actin fibers along the retractor muscle (’rm’). Vasa2+ cells are less abundant towards the oral and aboral ends of the mesentery, with only few Vasa2+ cells present in the physal region (E’’). (G-I) Confocal imaging stacks of cross sections of adult female gonadal mesentery immunostained for Vasa2 (green), tubulin (white), and EdU-labelled for S-phase (3 days pulse, red). Small, basiepithelial Vasa+ cells are found in the reticulate and gonad tracts (arrowheads), partially displaying EdU incorporation (asterisks). Some Vasa2+/EdU+ cells consist of duplets (I), which suggests that they recently divided. Blue: DAPI DNA dye. cgt: cnidoglandular tract; ct: ciliated tract; rt: reticulate tract; sg: somatic gonad; o: oocyte; rm: retractor muscle. Scale bars: 10μm (B-E’’’) and 50μm (H, I).

**Figure S4.**
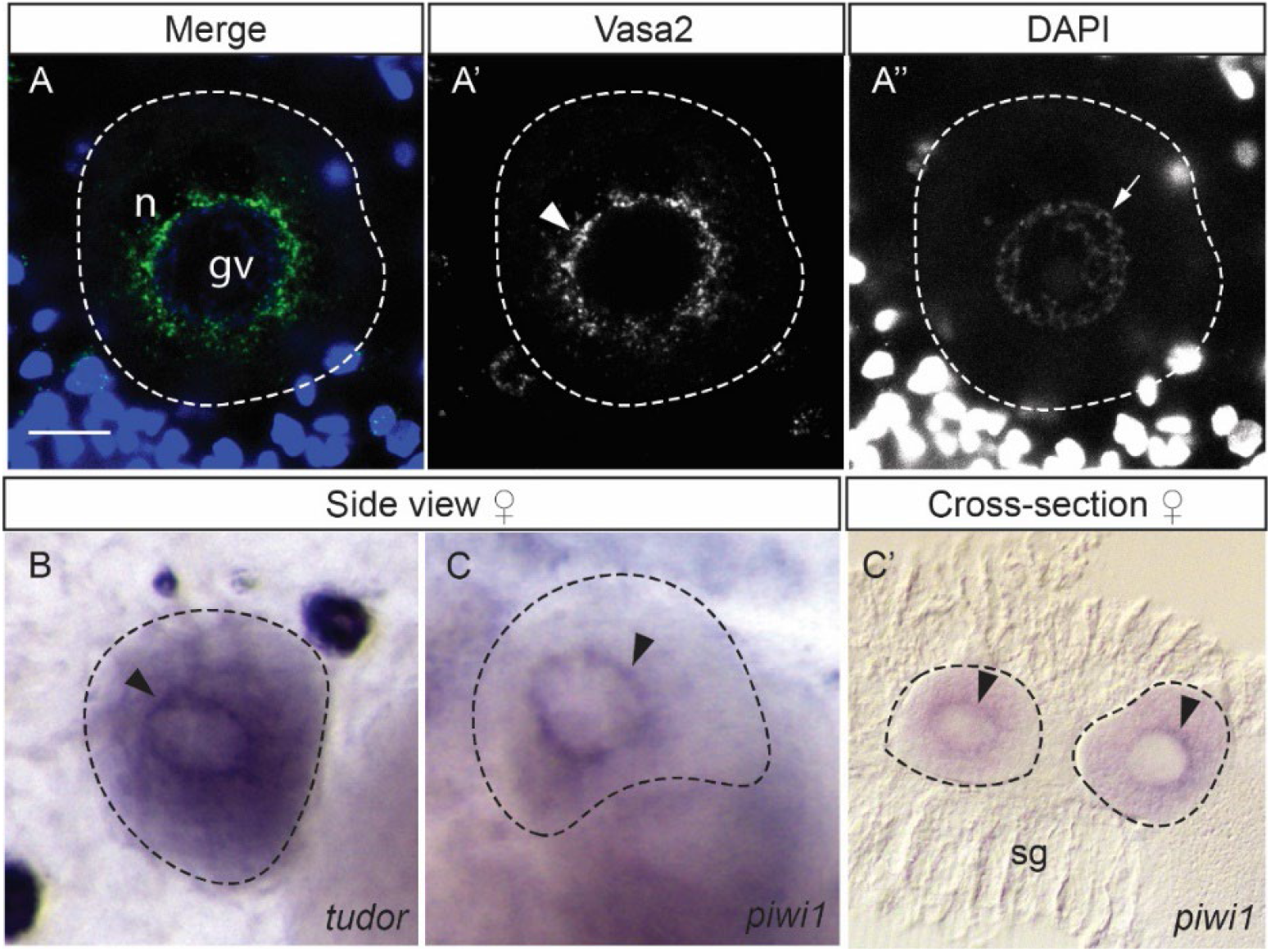
Evidence for the concentration of Vasa2 protein. (A-A’’), *tudor* (B) and *piwi1* mRNAs (C,C’) at perinuclear granules (*nuage*) of developing oocytes. (A-A’’) Confocal imaging stacks of a single oocyte, close up of Figure 3C. Vasa2 protein is detected in perinuclear granules (i.e. *nuage*) around the oocyte germinal vesicle (A, green; A’, arrowhead). DNA appears lowly condensed in the germinal vesicle (A, blue; A’’, arrow). (B-C’) ISH of *tudor* (B) and *piwi1* (C, C’) genes on gonadal tissue pieces viewed from lateral (B, C) or at a cross section (C’). B and C are close ups of Figure S1E and 1B, respectively. Arrowheads: nuage. Blue: Hoechst DNA dye. gv: germinal vesicle; n: nuage; sg: somatic gonad. Scale bar: 10μm.

**Figure S5.**
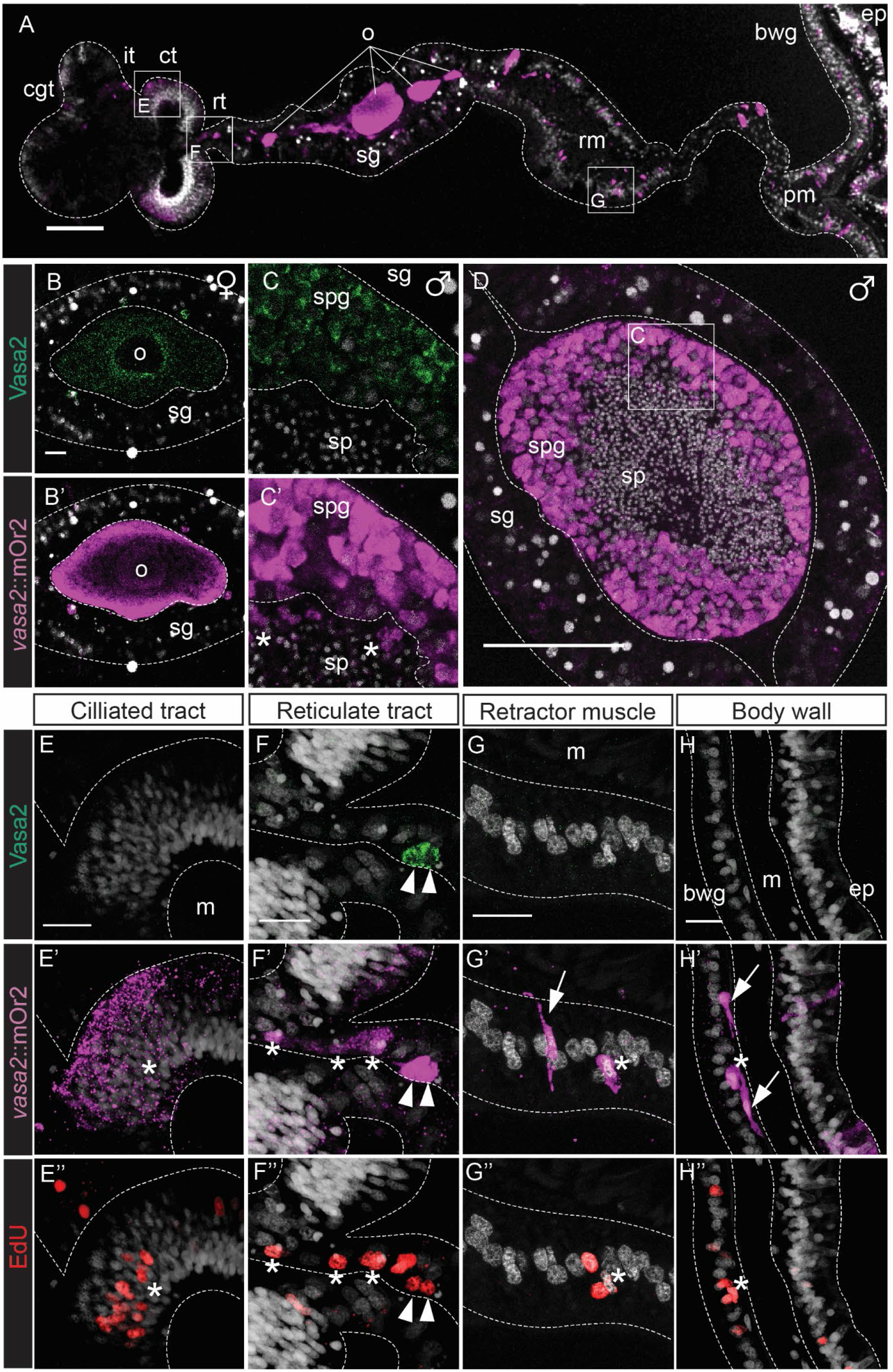
Expression of *vasa2* ∷mOr2 highlights germinal and somatic progeny of Vasa2+/Piwi1+ stem-like cells in the adult gastrodermis. (A-H) Confocal imaging stacks of cross sections of *vasa2∷mOr2* female and male mesenteries immunostained for Vasa2 (B, C, E-H, green), mOr2 (A, B’, C’, D, E’-H’, magenta) and labelled for S-phase by EdU (E’’-H’’, 3-days EdU pulse, red). In the gonadal tract, mOr2 colocalizes with Vasa2 to oocytes (A-B’, o) and spermatogonia (C-D, spg). mOr2 protein is likely transferred to differentiating, Vasa2-spermatocytes (C’, asterisks) by cytoplasmatic inheritance. (E-H’’) Close ups of specific mesentery regions (boxed in A) show high mOr2 levels in Vasa2+/EdU+ cells of the reticulate tract (F-F’’, arrowheads). In contrast, relatively low mOr2 levels are detected in Vasa2-/EdU+ (E-H’’, asterisks) and Vasa2-/EdU– (G’, H’ arrows) cells in the distal ciliated tract (E’-E’’), reticulate tract (F’-F’’), retractor muscle (G’-G’’) and body wall (H’-H’’). Grey: Hoechst DNA dye. cgt: cnidoglandular tract; it: intermediate tract; ct: ciliated tract; rt: reticulate tract; sg: somatic gonad, m: mesoglea; o: oocyte; spg: spermatogonia; sp: sperm; rm: retractor muscle; pm: parietal muscle; bwg: body wall gastrodermis; ep: epidermis. Scale bars: 50μm (A, D) and 10μm (B, C, E-H).

**Figure S6.**
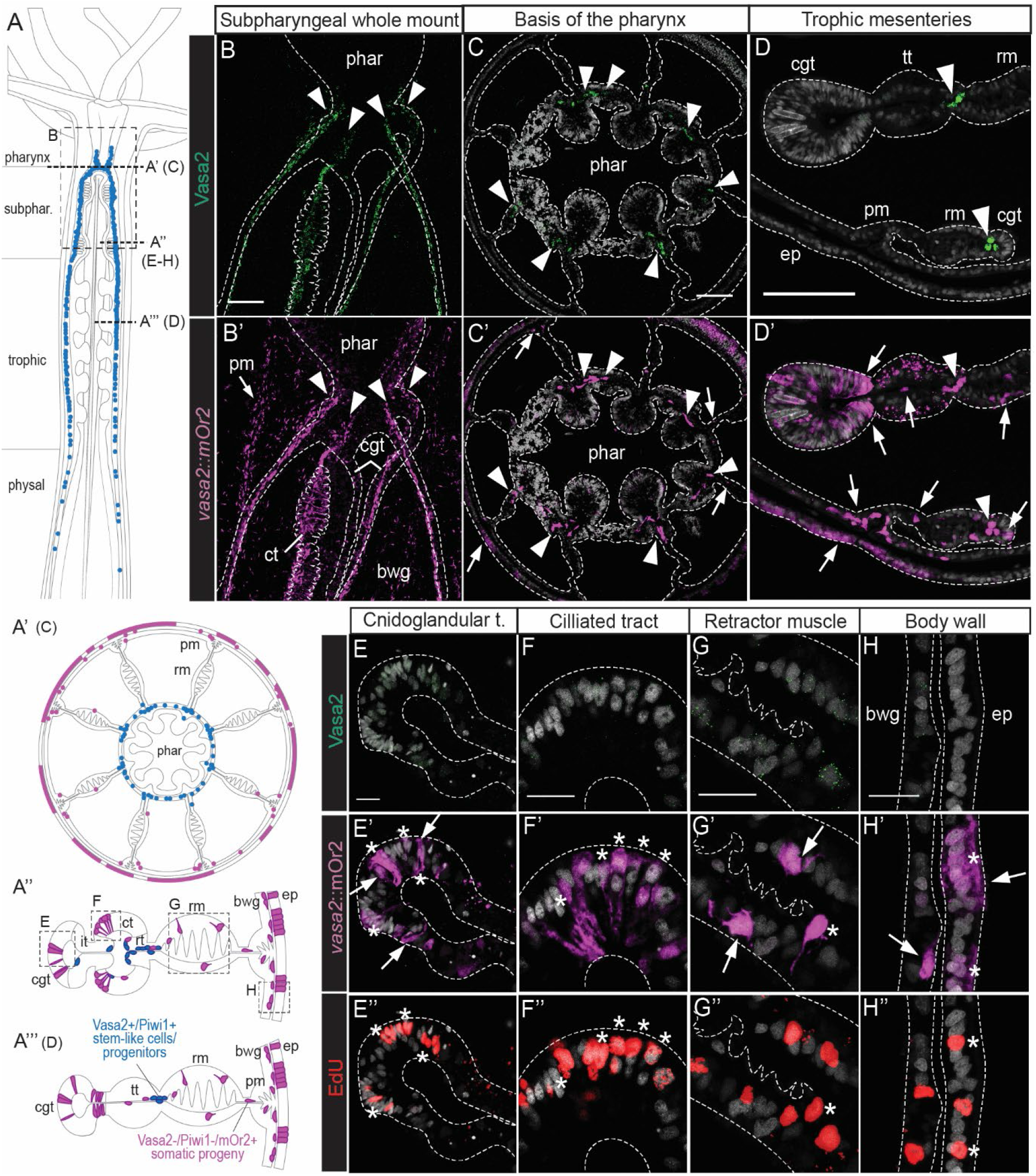
Expression of *vasa2* ∷mOr2 highlights abundant somatic progeny of mesenterial Vasa2+/Piwi1+ stem-like cells in the gastrodermis of growing juveniles. (A-A’’’) Schematics of longitudinal (A) and cross sections (A-A’’’) through pharyngeal (A’), subpharyngeal (A’’) and trophic (A’’’) regions. Blue: Vasa2+/mOr2+ cells; violet: Vasa2–/mOr2+ cells. (B-H’’) Confocal imaging stacks of a *vasa2∷mOr2* juvenile as whole-mounts (B-B’) or cross-sectioned (C-H’’) as indicated in (A-A’’’), immunostained for Vasa2 (B-H, green) and mOr2 (B’-H’, magenta), and labelled for S-phase (E’’-H’’, 1 day EdU pulse, red). Vasa2+/mOr2+ cells are detected between the septal filament and retractor muscle tract and extend oral-aborally from the basis of the pharynx (B-B’, C-C’, arrowheads) to the physal region (D-D’, arrowheads). Abundant Vasa2–/mOr2+ cells (B’-H’, arrows), which are often EdU+ (E’-H’’, asterisks), are found scattered throughout the gastrodermis. Grey: Hoechst DNA dye. phar: pharynx; cgt: cnidoglandular tract; it: intermediate tract; ct: ciliated tract; rt: reticulate tract; tt: trophic tract; rm: retractor muscle; pm: parietal muscle; bwg: body wall gastrodermis; ep: epidermis. Scale bars: 100μm (B-B’), 50μm (D, H-H’’) and 10μm (C, E-G’’).

**Figure S7.**
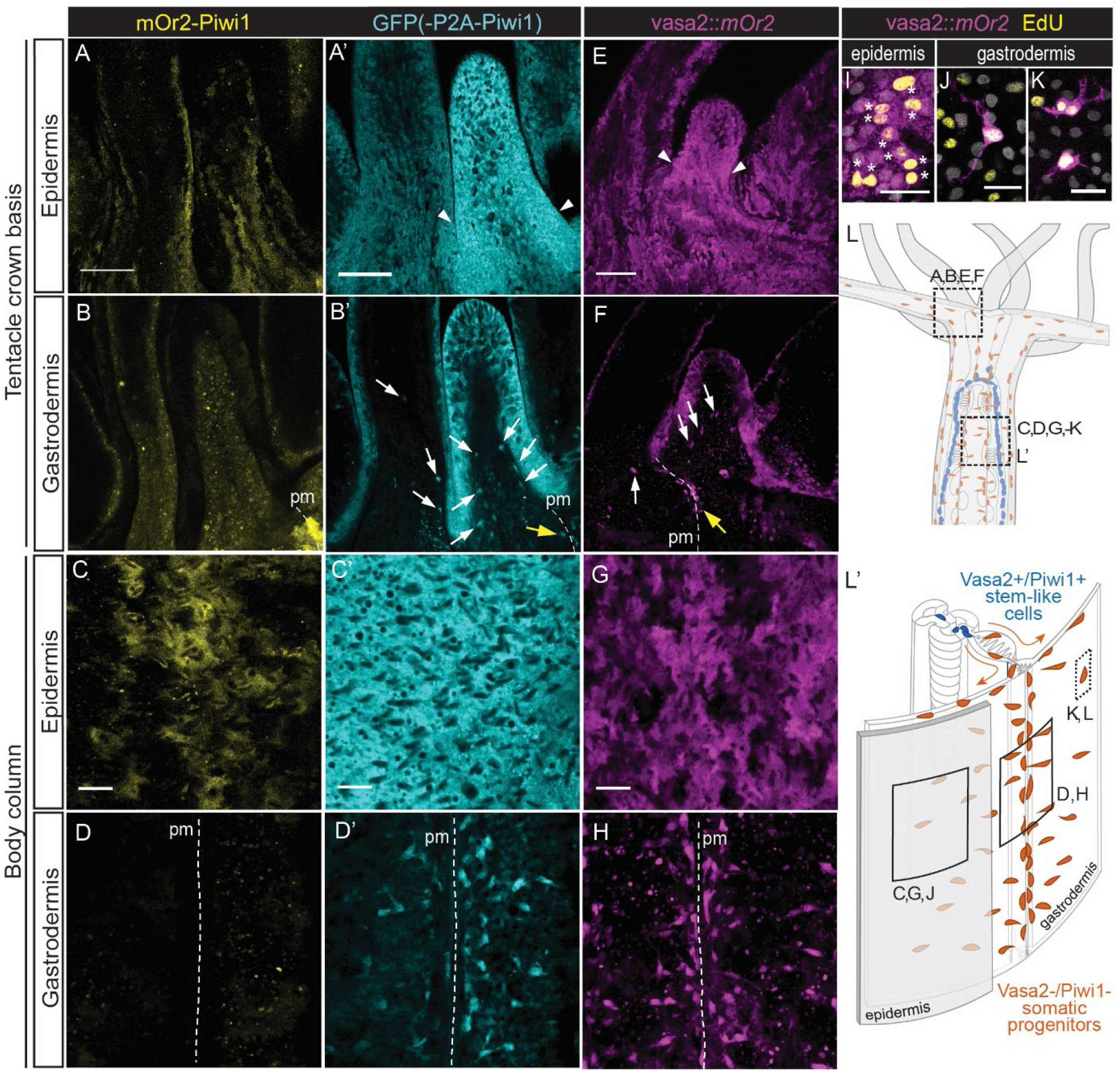
Characterization of the tentacle and body wall reporter expression in *piwi1^mOr2/P2A-GFP^* and *vasa2∷mOr2* polyps. (A-H) *In vivo* confocal imaging stacks of the epidermis and gastrodermis of whole mount *piwi1^mOr2/P2A-GFP^* (A-D’) and *vasa2∷mOr2* (E-H) juveniles at the basis of the tentacle crown and at the body wall as indicated in (L, L’). In the epidermis, mOr2-Piwi1 is found in patches of cells (A, C), while GFP appears present ubiquitously in all cells (A’, C’). *vasa2:*:mOr2 is expressed in patches of cells in the epidermis (E, G). Both vasa2∷mOr2 and GFP(-P2A-Piwi1) are found at higher levels in the epidermis of growing tentacles (A’, E’, arrowheads) in comparison to adjacent, developed ones. In the gastrodermis no specific mOr2-Piwi1 is detected (B, D) with the strong signal in the parietal muscle corresponding to endogenous red fluorescence (Ikmi and Gibson, 2010). Gastrodermal GFP+ and *vasa2:*:mOr2+ cells are concentrated along the parietal muscle tracts of the body column (D’, H) up to the basis of the tentacles (B’, F, yellow arrows) and are also found extending into the tentacle gastrodermis (B’, F, white arrows). (I-K) Confocal imaging stacks of a whole mount *vasa2∷mOr2* polyp immunostained for mOr2 (magenta) and EdU (1h pulse, yellow). Areas imaged as indicated in (L, L’). Pairs of cells with similar EdU levels after 1h pulse are found within mOr2+ epidermal cell patches (I, asterisks). mOr2+, neuron-shaped cells are found in the body wall gastrodermis (J, K), some forming duplets (K). (L-L’) Schematic of the oral half of a juvenile polyp (L) and a 3D illustration of a piece of body column (L’) highlighting the location of mesenterial Vasa2+/Piwi1+ stem-like cells (blue) and their likely migratory gastrodermal progeny cells (orange). Epidermis coloured in grey. pm: parietal muscle. Scale bars: 50μm in (A, A’, E) and 5μm in (C, C’, G, I-K).

**Figure S8.**
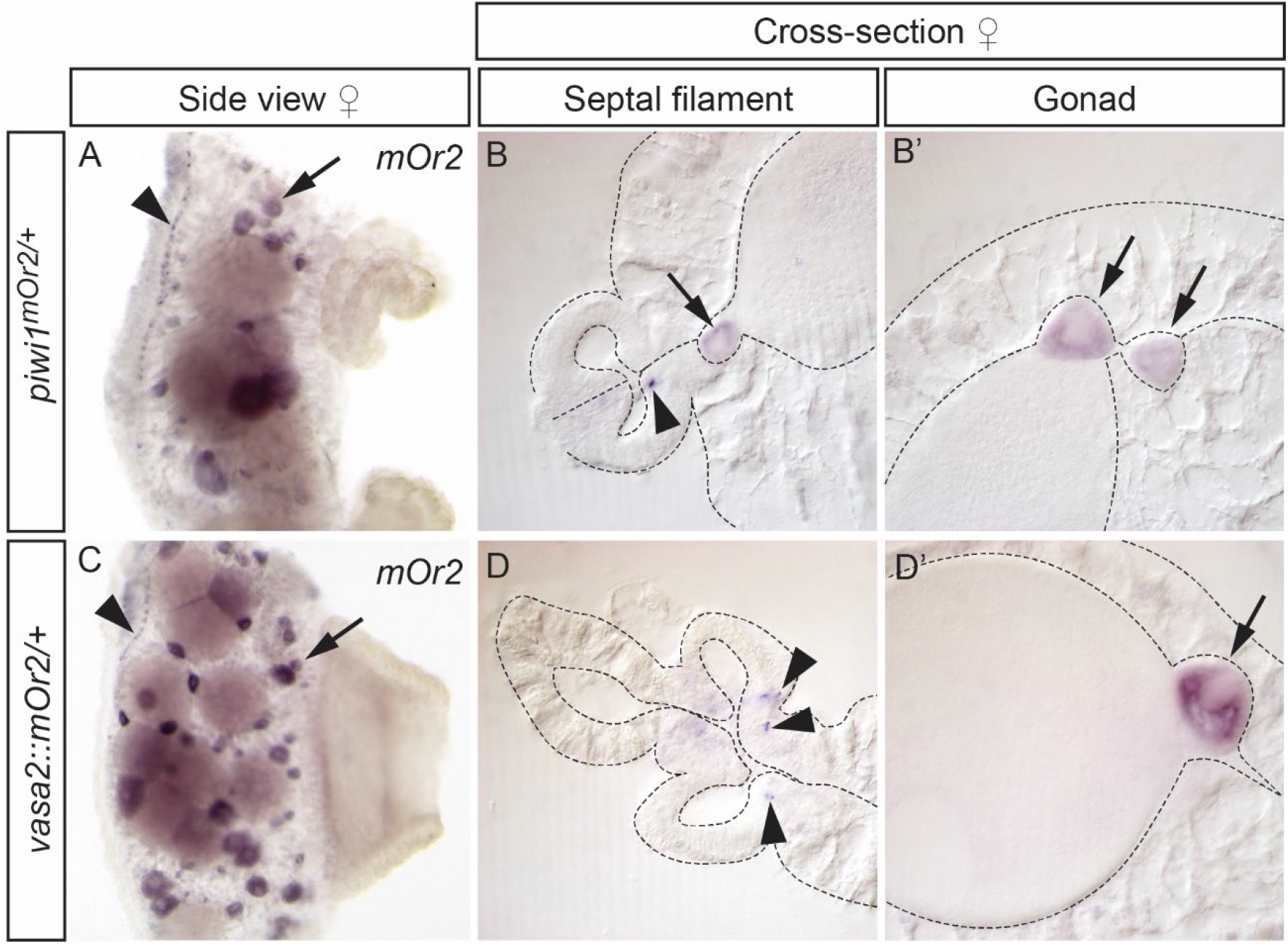
Detection of *mOr2* mRNA in *vasa2∷mOr2* and *piwi1^mOr2^* adult female gonad tissues. (A-D’) Whole mount ISH of the *mOr2* gene on female gonadal mesentery tissues of a *piwi1^mOr2/+^* (A-B’) and a *vasa2∷mOr2* (C-D’). In both lines, *mOr2* is only detected in small to medium oocytes (A, B, C, D’, arrows) and in basiepithelial cells along the reticulate tract (A, B, C, D, arrowheads), as found for *piwi1* mRNA (see Figure 1A-A’). (A, C): Side views of tissue pieces. (B, B’, D, D’): Cross sections.

**Figure S9.**
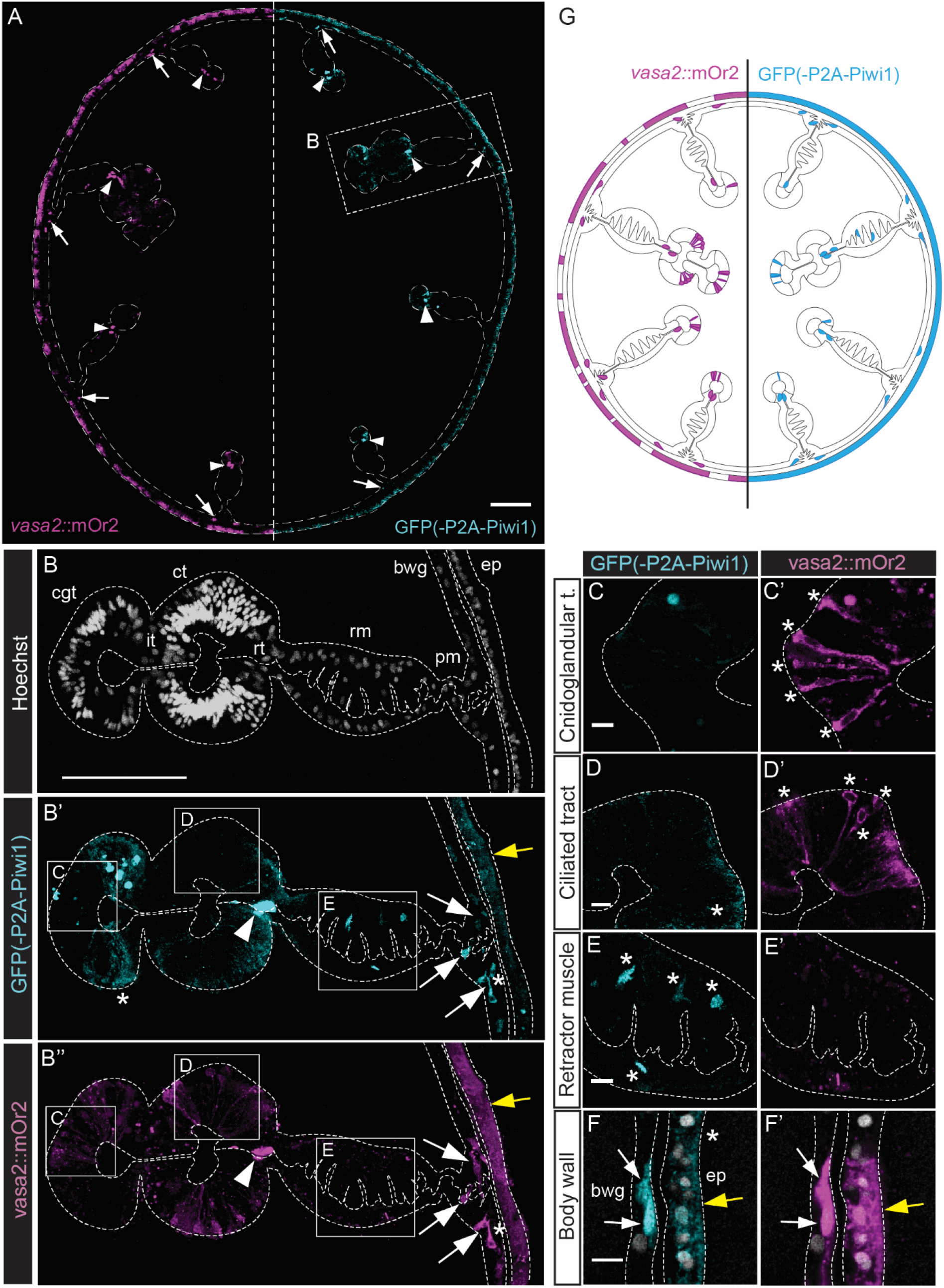
Partial colocalization of *vasa2* ∷mOr2 and GFP(-P2A-Piwi1) in juveniles. (A-F’) Confocal imaging stacks of juvenile subpharyngeal mesentery cross sections expressing both *piwi1^P2A-GFP^* and *vasa2∷mOr2* immunostained against GFP (cyan) and mOr2 (magenta). Cross section of the body column (A), a full mesentery (B-B’’), the mesenterial tracts indicated in B’ and B’’(C-E’), and body wall (E-E’). High mOr2 and GFP levels colocalize to stem-like cells at the basis of the septal filament, adjacently to the retractor muscle (A, B’, B’’, arrowheads). Lower levels of mOr2 and GFP colocalize to cells around each of the parietal muscle tracts (A, B’, B’’, white arrows), the body wall gastrodermis (F, F’, white arrows) and epidermis (B’, B’’, F, F’, yellow arrows). No colocalization is found in GFP+/mOr2-cells in the cnidoglandular and parietal muscle tracts (B’, asterisks), ciliated tract (D, asterisk), retractor muscle tract (E, asterisks) and epidermis (G, asterisk). GFP-/mOr2+ cells are found in cnidoglandular tract (C’, asterisks) and ciliated tract cells (D’, asterisks). (G) Schematic of a polyp cross section depicting the reporter line expression patterns observed for *vasa2:*:mOr2 and GFP(-P2A-Piwi1) as in A-F’. Grey: Hoechst DNA dye. cgt: cnidoglandular tract; it: intermediate tract; ct: ciliated tract; rt: reticulate tract; rm: retractor muscle; pm: parietal muscle; bwg: body wall gastrodermis; ep: epidermis. Scale bars: 50μm (A, B) and 5μm (C-F).

